# Super-delta2: An Enhanced Differential Expression Analysis Procedure for Multi-Group Comparisons of RNA-seq Data

**DOI:** 10.1101/2021.01.30.428977

**Authors:** Zihan Cui, Yuhang Liu, Jinfeng Zhang, Xing Qiu

## Abstract

**Background:** We developed `super-delta2`, a differential gene expression analysis pipeline designed for multi-group comparisons for RNA-seq data. It includes a customized one-way ANOVA F-test and a post-hoc test for pairwise group comparisons; both are designed to work with a multivariate normalization procedure to reduce technical noise. It also includes a trimming procedure with bias-correction to obtain robust and approximately unbiased summary statistics used in these tests. We demonstrated the asymptotic applicability of `super-delta2` to log-transformed read counts in RNA-seq data by large sample theory based on Negative Binomial Poisson (NBP) distribution.

**Results:** We compared `super-delta2` with three commonly used RNA-seq data analysis methods: limma/voom, edgeR, and DESeq2 using both simulated and real datasets. In all three simulation settings, `super-delta2` not only achieved the best overall statistical power, but also was the only method that controlled type I error at the nominal level. When applied to a breast cancer dataset to identify differential expression pattern associated with multiple pathologic stages, `super-delta2` selected more enriched pathways than other methods, which are directly linked to the underlying biological condition (breast cancer).

**Conclusions:** By incorporating trimming and bias-correction in the normalization step, `super-delta2` was able to achieve tight control of type I error. Because the hypothesis tests are based on asymptotic normal approximation of the NBP distribution, `super-delta2` does not require computationally expensive iterative optimization procedures used by methods such as edgeR and DESeq2, which occasionally have convergence issues.

## Introduction

High-throughput technologies for measuring gene expressions have become an indispensable component of modern biomedical research in recent years. Due to the high-dimensional nature of whole transcriptome gene expression profiles, most researchers need to select a subset of “interesting genes” to answer their questions at hand. This is typically achieved via differential gene expression analysis (DGEA) that identifies genes with significantly different mean expression levels across two or more phenotypic groups. It is well-known that various types of systematic noises exist in high-throughput gene expression data, thus it is a common practice to apply normalization procedures [1–6] to reduce these noises to enhance the DGEA. Most popular normalization procedures are data-driven transformations designed to ensure certain statistical characteristics are constant across all transformed samples. For example, the global normalization [1], also known as reads per million (RPM) or counts per million (CPM) normalization for RNA-seq data, is popular for both microarray and RNA-seq data. A trimmed variant of global normalization called TMM [4] is also widely used in RNA-seq data analysis. After this normalization, all samples are guaranteed to have constant (trimmed) mean expression levels. The median-IQR normalization [2] ensures both median and IQR of expressions of every sample are the same. Motivated by the quantile-quantile plot, the quantile normalization [3] is designed to equalize all quantiles (or equivalently, the empirical distribution function) of the samples. Although normalization procedures can reduce the variability in the raw data and make the DGEA more stable, this variance reduction is typically accompanied with certain bias because these procedures have to borrow information from all genes, including both non-differentially expressed and differentially expressed genes. This bias is most obvious when the data exhibits unbalanced differential pattern, namely, there are significantly more up-regulated genes than down-regulated genes or *vice versa*, and it can reduce statistical power and/or inflate type I error, especially when the sample size is relatively large [7, 8].

Recently, we developed a new differential expression analysis pipeline, dubbed as `super-delta` [9]. This method consists of three conceptual components: a) a multivariate extension of the global normalization to reduce technical noise; b) a robust trimming procedure designed to minimize the bias introduced by the normalization step; and c) a modified t-test optimized for the first two steps which is asymptotically unbiased based on theoretical derivations. Together, a) and b) serve as a *robust normalization method* for the last step. Using extensive simulations, we showed that `super-delta` achieved the best overall statistical power with tight control of type I error rate than its competitors, and the performance of `super-delta` was close to that of an *oracle test*, defined as the two-sample t-test applied to simulated data without technical noise. We also applied `super-delta` and other methods to a set of microarray data collected from breast cancer patients who responded differently towards neoadjuvant chemotherapy. `super-delta` was able to identify more differentially expressed genes (DEGs) than its competitors, and these DEGs are more biologically relevant to chemotherapy responses of breast cancer than those selected by the other methods.

Despite those advantages, `super-delta` still has two main weaknesses. First, it only applies to two-group comparison therefore is not capable of performing one-way ANOVA F-test for multi-group comparisons. Note that unlike most other normalization methods, a *modified* t-test must be used to take the full advantage of the robust normalization method implemented in `super-delta`. Therefore, it is not trivial to extend `super-delta` from two-group comparison to one-way ANOVA. Secondly, the particular form of the modified t-test was derived by theoretical derivations based on a *normal* mixed effects model. It is not immediately clear whether `super-delta` is applicable for RNA-seq data, which are typically modeled by discrete distributions such as negative binomial Poisson (NBP) distributions[10].

In this study, we proposed to extend `super-delta` so that it can be used for multi-group comparisons. Dubbed as `super-delta2`, it has: 1) a one-way ANOVA F-test that uses a combination of multivariate extension of the global normalization method and robust trimming to remove the impact of technical noise to the F-statistic; 2) a *post-hoc* pairwise group comparison method that is an extension of the Tukey’s method optimized for the same robust normalization procedure. In addition, we performed theoretical analyses based on the NBP distribution and showed that the modified F-test and Tukey’s test statistics are asymptotically normal, therefore `super-delta2` is valid for RNA-seq data. We designed thorough simulation studies based on the NBP distribution and compared `super-delta2` with three widely used RNA-seq analysis pipelines: limma/voom[11, 12], edgeR [13], and DESeq2[14]. Overall, `super-delta2` achieved the best balance between type I error and statistical power; and it was the only method that controlled type I error at the nominal level in all simulation settings. The advantage of `super-delta2` is also evident when we applied it and competing methods to a large collection of breast cancer gene expression data, which are divided into three groups according to the pathologic stages of the samples. Overall, `super-delta2` selected more biologically meaningful DEGs, as shown by subsequent pathway analyses.

## Materials and Methods

### A mixed-effects model for super-delta2

In this study, we model the log2-transformed gene expressions by the following mixed effects model, which is a multi-group extension of a similar model used by the original `super-delta` method [9] and many other studies [7, 8, 15–18].

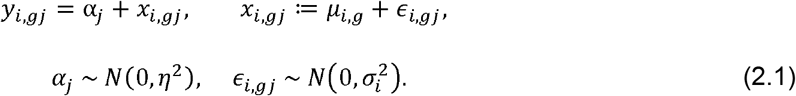

Here *i* = 1,2,…, *m* is the gene index, *g* = 1,2,…, *G* is the group index, and *j* = 1,2,…, *N*_*g*_ is the sample index. *y*_*i,gj*_, the observed gene expression, is decomposed into several parts: a) α_*j*_, a random effect term that models sample-specific variations commonly known as the technical noise, and b) *x*_*i,gj*_, the *oracle gene expression* that is free of technical noise, which is further divided into *μ*_*i,g*_ (per-group mean expression levels) and ε_*i,gj*_, which represents both meaningful biological variation and the *i.i.d.* measurement error. Most normalization procedures strive to recover *x*_*i,gj*_ from *y*_*i,gj*_ with the price of a small bias [7], then apply a standard DGEA such Welch t-test to 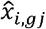 (the normalized gene expressions) to obtain *p*-values. As an alternative approach, the original `super-delta` method [9] uses a three-step algorithm to obtain *p*-values without the explicit estimation (normalization) of 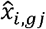 : 1. We compute *δ*_*ik,·*_ (the “deltas”), the pairwise differences between genes *i* and *k*. According to Model (2.1), sample-specific variation (α_*j*_) is removed by this step. 2. A modified t-test is applied to these deltas to obtain *t*_*ik*_, for all *i,k*, = 1,2,…, *m*. 3. Finally, a robust median fold trim median (MFTM) estimator is used to summarize those *t*_*ik*_ into one representative t-statistic (denoted by 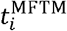), based on which the *p*-value is computed.

As an extension to the original `super-delta, super-delta2` also removes sample-specific variation by the *δ*-step. To handle multi-group cases, we have to use a completely different trimming strategy that works with a modified one-way ANOVA F-test in `super-delta2`. We provide technical details of `super-delta2` in the following sub-sections.

### The delta-step and the oracle F-statistics

First, we compute the “deltas” according to the following formula

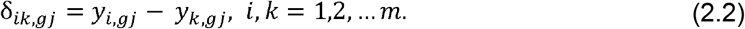

Based on Equation (2.1), 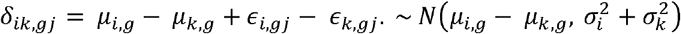. Note that sample-specific technical noise (*α*_*j*_) is cancelled in these deltas.

Assume that the oracle data, *x*_*i,gj*_ ≔ *μ*_*i,g*_ + *ϵ*_*i,gj*_, are available to us and we would like to test the following hypotheses for multi-group comparisons

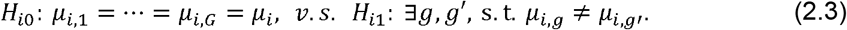

The corresponding one-way ANOVA F-test can be expressed as follows

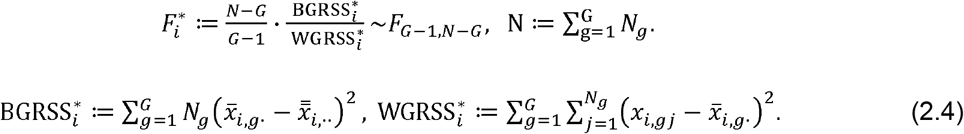

In the above formula, 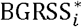 is between-group residual sum of squares and 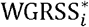 is within-group residual sum of squares. Unfortunately, because they are both computed from the oracle data instead of the actual observations (*y*_*i,gj*_), Equation (2.4) is not directly applicable in practice. We will describe fast and efficient ways to estimate 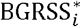 and 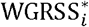 from the observations in the next two subsections.

### Estimation of Within-Group Residual Sum of Squares

By exploiting the fact that within-group variation is invariant under per-group shift transformation, we propose an efficient algorithm to estimate 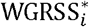 based on global normalization. Let 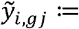 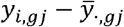 be the globally normalized gene expressions; let 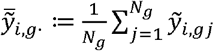 be the within-group mean of 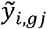, and 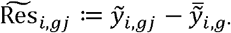 be the residual of globally normalized gene expressions within group *g*. Let 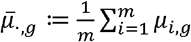 be the mean of the expected gene expression levels in group *g*; 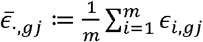 be the mean variation pertain to the *j*th sample in group *g* averaged over all genes; and 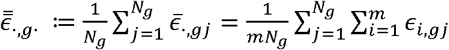 be the average variations for all genes and samples in group *g*. Putting all these notations together, we have

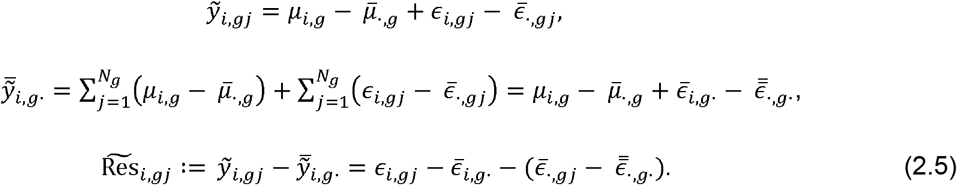

Note that the mean values 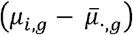 get canceled out in 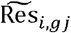, and the second term, 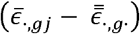, is of order *O*(*m*^−1/2^max_*i*_*σ*_*i*_). So we have 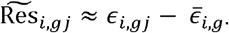 under an additional assumption that

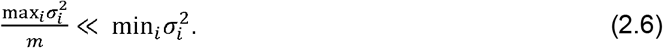

In other words, per-gene variance terms 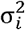 are not so different, to the point that the largest 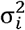 is of order *m*, times larger than the smallest 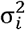. We believe this assumption is reasonable for real data, especially after the non-specific filtering of low read count genes.

Recall that

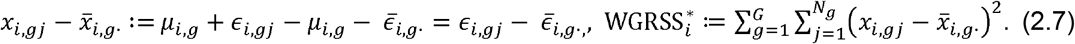

By replacing 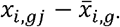 with 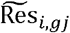, the estimates of 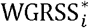 and per-gene variance 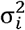, are

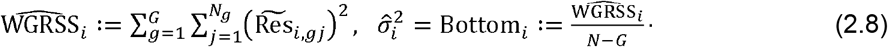

### Estimation of Between-Group Residual Sum of Squares

The estimation of BGRSS requires more careful considerations, because it is not invariant to per-group location transformations. Recall that the BGRSS to be estimated is:

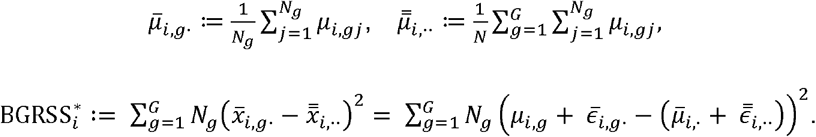

On the other hand, the between-group residuals computed from the deltas, denoted by *R*_*ik,g*_, are

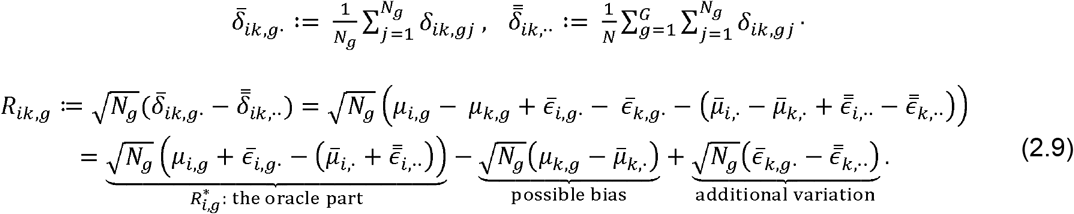

In the above equation, 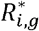 (the oracle part) is what we are after because 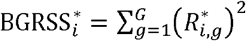. Let 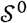 and 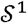 be the indices of non-differentially expressed genes (NDEGs) and differentially expressed genes (DEGs), respectively. The possible bias term equals zero for 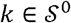, because 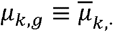 under null hypotheses. It is nonzero for 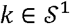 therefore may skew the F-test. The last term has variance 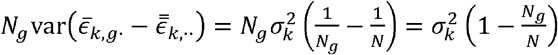. Imagine that we could use an *oracle trimming* method to remove exactly all 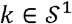 and take the trimmed mean of *R*_*ik,g*_ in the direction of *k*. We can define an estimate of 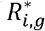 as follows

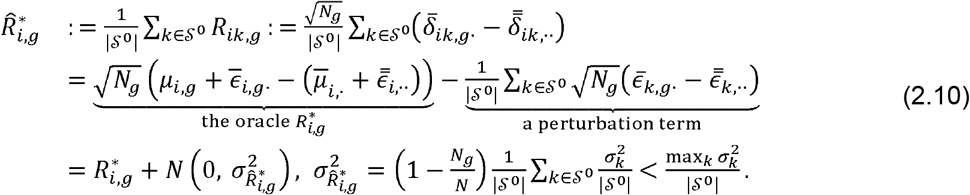

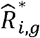 converges to 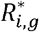 very fast because 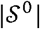, the number of NDEGs, is typically very large. In practice, the oracle information is not available, so we propose the following spherically trimmed mean estimator for 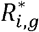 and 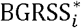. Let 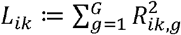 be the squared Euclidean length of vector *R*_*ik,·*_. For a pre-specified *q* ∈ (0,1), let 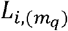, *m*_*q*_ ≔ [(*m* − 1)(1 − *q*)], be the (1 − *q*)th empirical quantile of *L*_*i·*_. We define an index set 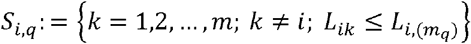, and the trimmed BGRSS estimator as

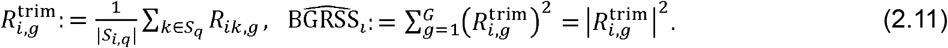

In other words, genes with extremely large between-group RSS are removed from the estimation of BGRSS. We noted that when *q*, the trimming proportion, is large (e.g., *q* > 0.2), this trimming procedure may under-estimate BGRSS and reduce the statistical power of `super-delta2`. We developed a bias-correction procedure based on moment-matching to reduce this bias. See Appendix 1 in Supplementary Text for more details.

Finally, the oracle WGRSS and BGRSS in Equation (2.4) are substituted by 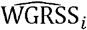 and 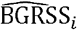 respectively to obtain the modified F-statistics and the corresponding *p*-values.

### Post-hoc Analysis in Tukey’s Style

In most multi-group comparisons, practitioners are also interested in post-hoc pairwise comparisons, to answer more specific questions such as which two clinical groups have significantly different mean read counts. We develop a Tukey’s style post-hoc pairwise test for `super-delta2` in this subsection. Recall that Tukey’s post-hoc test can be formulated as follows

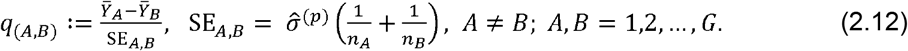

Here 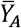 and 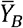 are the means of groups A and B; SE is the standard error of 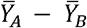; and 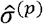 is the pooled standard deviation calculated from observations in *all groups*.

In our case, we propose to replace 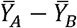 by 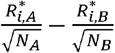, because

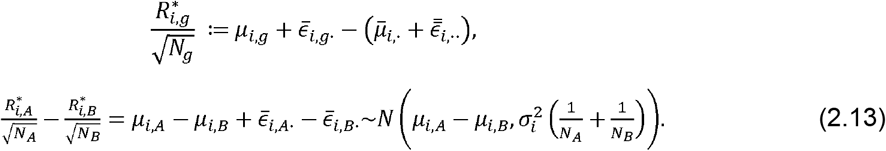

Now we replace 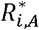 and 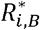 by their trimmed estimates; 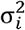 by 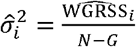 defined by Equation (2.8); and define

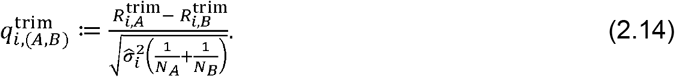

Based on the construction, 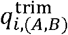 approximately follows t-distribution *T*(*N* − *G*) if *μ*_*i,A*_ − *μ*_*i,B*_ = 0, based on which we can obtain *p*-values for pairwise group comparisons. Note that the degree of freedom is *N* – *G* instead of *N*_*A*_ – *N*_*B*_ – 2 in two-sample t-test, because 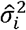 is a *pooled* variance. This is a main advantage of Tukey-style post-hoc analyses compared with the simpler approach that directly apply two-sample t-test to all pairwise comparisons.

A complete description of the `super-delta2` algorithm is provided in Appendix 3 in Supplementary Text.

### Applicability of super-delta2 to RNA-seq Data Based on Negative Binomial Poisson Models

Unlike microarray data, raw RNA-seq data are represented as *integer-valued* read counts that are best modeled by an appropriate discrete distribution instead of normal distribution, especially for those genes with very low mean counts thus more granularity. By and large, two approaches exist for conducting DGEA for RNA-seq data. In the first approach [10, 13, 14, 19, 20], the read counts are modeled by a discrete distribution such as Poisson distribution and correlated with covariates, such as binary group labels for multi-group comparisons, with the corresponding generalized linear model (GLM). P-values are obtained by the likelihood ratio test or Wald test for the GLM. Originally, Poisson model and Poisson GLM was used for this purpose [20], but later it was shown that real RNA-seq data exhibits extra-Poisson variation (i.e., overdispersion) which can be accounted for by negative binomial Poisson (NBP) model [10]. We want to point out that the small sample null distribution of the test statistics in likelihood ratio or Wald tests for those GLMs are unknown therefore we can only obtain *approximate* p-values based on large sample approximation (usually based on *χ*^2^-distributions).

In the second approach [12, 21, 22], a combination of non-specific filtering (to remove those genes with extremely low read counts and large granularity), variance-stabilization transformation (to reduce the skewness of the data), and normalization (to reduce technical noise) is used to make the transformed distribution closer to normal distributions; then a Central Limit Theorem (CLT) is applied to justify that a standard linear regression model and the associated F- and t-tests are asymptotically valid when the sample size is relatively large. Although both approaches rely on large sample approximations, the second approach tended to perform slightly better than the first approach in comparative studies [21, 22]. We think two reasons may explain the advantage of the second approach: 1. Statistical methods developed for standard regression models are more mature than those developed for GLMs, and there is no need to rely on iterative optimization procedures so the algorithms are much faster and the convergence to global MLE is guaranteed by the Gauss-Markov theorem in the second approach. 2. Most data normalization methods turn the discrete distribution into a pseudo continuous distribution therefore are not compatible with the first approach. This fact greatly limits the ability of the first approach to reduce technical noise.

Given these considerations, we decide to follow the second approach and use NBP models in our simulation studies to compare the performance of `super-delta2` and competing methods. Note that while Model (2.1) is based on the normality assumption, most of the theoretical derivations in that section are asymptotically valid as long as 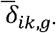, the “building blocks” for 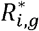, are asymptotically normal. Theoretical derivations on the asymptotic normality of 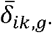 based on an NBP model for `super-delta2` is given in Appendix 2 in Supplementary Text.

### Brief Summary of Other Differential Gene Expression Analysis Methods

In this study, we compare the performance of `super-delta2` with three commonly used differential gene expression analysis methods on RNA-seq data: limma+voom, edgeR and DESeq2. We briefly introduce these methods as follows.

#### 1. limma+voom

Limma [11] was originally developed for differential expression analysis of microarray data. Voom [12] is an adaptation of Limma that is suitable for RNA-seq data. It generates a precision weight for each observation by estimating the mean-variance relationship of the log-counts and enters these weights into the limma empirical Bayes analysis pipeline. Together they allow flexible and powerful analyses of gene expression data.

#### 2. edgeR

edgeR[13] implements novel statistical methods based on negative binomial distribution as a model for count variability, including empirical Bayes methods, exact tests, and generalized linear models. edgeR is especially suitable for analyzing designed experiments with multiple experimental factors but possibly small numbers of replicates.

#### 3. DESeq2

DESeq2[14] performs an internal normalization in order to correct RNA composition bias at first, then uses shrinkage estimations for dispersions and fold changes. It can detect outliers using Cook’s distance and remove these genes from analysis. DESeq2 fits negative binomial generalized linear models for each gene and uses the Wald test for significance testing.

## Simulation Studies

An NBP distribution is an integer-valued distribution with three parameters, the location parameter *v*, and two shape parameters *κ* and *a*. The mean and variance of *X* ~ NBP(*v,κ,a*) are

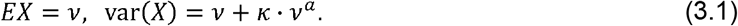

Other technical details of the NBP distribution, such as the probability density function and its relationship with the negative binomial distribution are provided in Appendix 2 in Supplementary Text. Since Equation (3.1) describes a nonlinear relationship between the mean and variance of genes, we can apply nonlinear regression on the sample mean and variance of read counts of real data to obtain rough estimates of these parameters. Specifically, two sets of real RNA-seq data were used for this purpose. The first dataset is a liver cancer (LIHC) RNA-seq data with n=423 samples provided by The Cancer Genome Atlas (TCGA, https://cancergenome.nih.gov/). A second dataset contains 1,212 breast cancer (BRCA) samples, also obtained from TCGA. More details about them are provided in section “Real Data Analysis”. The empirically estimated nonlinear relationship between means and variances of gene expressions in these two datasets are summarized in Figure 1. We see that parameter *a*, which is either 2.2 or 2.0, is more stable than *κ*, which seems to have a more flexible range.

**Figure 1:**
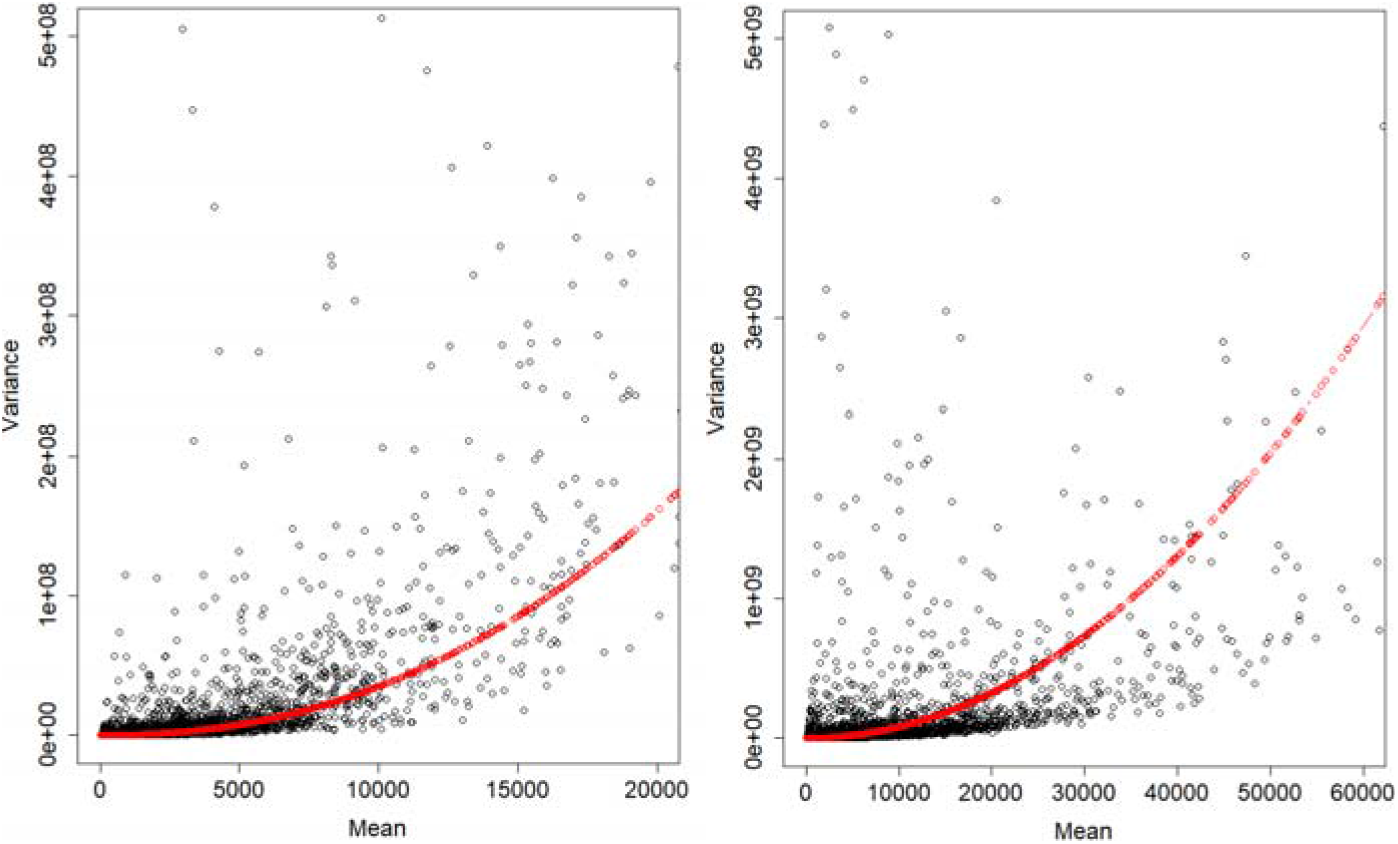
Scatterplot of individual genes’ mean values and variances in liver cancer (left) and breast cancer (right) and the regression line inferred from each sample. In liver cancer case, the regression function is variance = mean + 0.0676 * mean^2.2^, in breast cancer case, the regression function is variance = mean + 0.6899 * mean^2^.

Three simulation studies based on NBP with different modeling parameters were conducted to evaluate the performance of `super-delta2` and three DGEA methods commonly used for RNA-seq data: limma+voom, DESeq2, and edgeR. Note that `super-delta2` was applied to log2-transformed read counts; limma+voom has its own built-in variance-stabilization transformation; DESeq2 and edgeR were designed to work with read counts directly. In each scenario, we generate read counts of *m*, = 5,000 genes in three groups (Group A, B, C), each with *n*_*g*_ = 50 subjects. Sample specific noise, denoted by *α*_*gj*_, can alter the observed read counts by two different mechanisms that will be made clear later. For simplicity, *α*_*gj*_ in all three simulations are assumed to be uniformly distributed with lower bound *l* and upper bound *u*.

All four methods were used to test one-way ANOVA hypotheses, namely, whether each gene has the same expectation across all three groups. In addition, we also performed post-hoc pairwise group comparisons for each approach to the simulated data. Type I error (specifically, per-family error rate) and statistical power for both one-way ANOVA tests and post-hoc pairwise group comparisons were calculated from 100 repetitions. Receiver operating characteristic (ROC) curves were plotted to visualize the statistical performance of these methods. To avoid the complication of multiple testing adjustment in comparing the performance of different methods, DEGs were based on unadjusted p-values with significance level α < 0.05.

### Simulation 1

Let *Y*_*i,gj*_ be the read count of the th gene for sample *j* in group *g*. In simulation 1, *Y*_*i,gj*_ is generated from the following NBP model

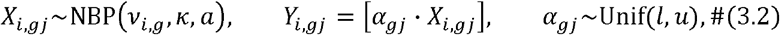

Here *κ* = 0.06 and *a* = 2.2 are two shape parameters in an NBP distribution, *X*_*i,gj*_ represent “noise-free” read counts, *α*_*gj*_ is a multiplicative technical noise for the *j* th sample in group *g*, [*x*] stands for the integer rounding function, *l* = 12, *u* = 30 are the lower and upper bounds of a uniform distribution. *v*_*i,g*_ are mathematical expectations of *X*_*i,gj*_ with the following values:

1. Group A was set as the baseline and *v*_*i,g*_ = 100, so that the expected mean counts of all genes are *EY*_*i,gj*_ ≈ *Eα*_*gj*_ · *v*_*i,A*_ = 2100.
2. For Group B, genes 1-600 are up-regulated with *v*_*i,B*_ = 150 so that *EY*_*i,gj*_ ≈ *Eα*_*gj*_ · *v*_*i,B*_ = 3150. Other genes have the same mean counts as group A (*v*_*i,B*_ = 100, *i* = 601,…5000).
3. For Group C, genes 401-1000 are down-regulated with *v*_*i,C*_ = 75 and *EY*_*i,gj*_ ≈ 1575. Other genes have the same mean counts as group A.

For each method, the average type I error rate, statistical power, and AUC value of simulation 1 are summarized in Table 1. By construction, the effect size (the difference of mean expression levels) is larger between Groups A and B than that between Groups A and C. Consequently, we have more power to detect DEGs when comparing A vs. B than comparing A vs. C. The comparison between Groups B and C is more complicated. Compared with the baseline (Group A): (a) genes 1 – 400 are up-regulated in Group B with a large effect size, (b) genes 401 – 600 are up-regulated in Group B *and* down-regulated in Group C with an even larger effect size, and (c) genes 601 – 1,000 are down-regulated in Group C with a small effect size. The expected statistical power for comparing B vs. C is therefore a weighted average of these cases. Based on Table 1, we found that `super-delta2` is more powerful than other methods, especially in the case B vs. C, which has both up- and down-regulated genes. What’s more, `super-delta2` is the only method that controlled type I error at the nominal significance level *α* = 0.05 for all tests.

**Table 1:** Type I error rate, statistical power, and AUC value of multi-group comparisons at significance level *α* = 0.05 for various methods in **simulation 1**. The 3^rd^ column (Overall) records statistical performance of the one-way ANOVA test, the rest three columns record results from post-hoc pairwise group comparisons. All reported results are averaged over 100 repetitions. (·) represents the standard deviation of these 100 repetitions.

|  | Method | Overall | A vs. B | A vs. C | B vs. C |
| --- | --- | --- | --- | --- | --- |
| Type I error | *super-delta2* | **0.048**<br>**(0.002)** | **0.050**<br>**(0.002)** | **0.048**<br>**(0.003)** | **0.048**<br>**(0.002)** |
|  | limma+voom | 0.145<br>(0.008) | 0.106<br>(0.006) | 0.065<br>(0.004) | 0.184<br>(0.009) |
|  | edgeR | 0.117<br>(0.004) | 0.079<br>(0.003) | 0.064<br>(0.004) | 0.152<br>(0.006) |
|  | DESeq2 | 0.131<br>(0.005) | 0.086<br>(0.004) | 0.071<br>(0.004) | 0.166<br>(0.008) |
| Statistical power | *super-delta2* | **0.974**<br>**(0.004)** | **0.996**<br>**(0.001)** | **0.900**<br>**(0.006)** | **0.963**<br>**(0.004)** |
|  | limma+voom | 0.921<br>(0.005) | 0.982<br>(0.002) | 0.858<br>(0.008) | 0.842<br>(0.008) |
|  | edgeR | 0.943<br>(0.005) | 0.993<br>(0.001) | 0.882<br>(0.007) | 0.888<br>(0.007) |
|  | DESeq2 | 0.944<br>(0.005) | 0.993<br>(0.001) | 0.884<br>(0.007) | 0.890<br>(0.006) |
| AUC<br>value | super-delta2 | **0.994**<br>**(0.001)** | **0.999**<br>**(0.000)** | **0.978**<br>**(0.002)** | **0.991**<br>**(0.001)** |
|  | limma+voom | 0.963<br>(0.003) | 0.990<br>(0.001) | 0.962<br>(0.003) | 0.912<br>(0.004) |
|  | edgeR | 0.978<br>(0.002) | 0.997<br>(0.000) | 0.968<br>(0.003) | 0.945<br>(0.004) |
|  | DESeq2 | 0.975<br>(0.002) | 0.997<br>(0.000) | 0.967<br>(0.003) | 0.941<br>(0.003) |

ROC curves of those four methods in simulation 1 were summarized in Figure 2. It shows that `super-delta2` has the highest AUC values in overall comparison. Once again, in comparison between Group B and Group C, the AUC value of `super-delta2` is significantly higher than the other three methods.

**Figure 2:**
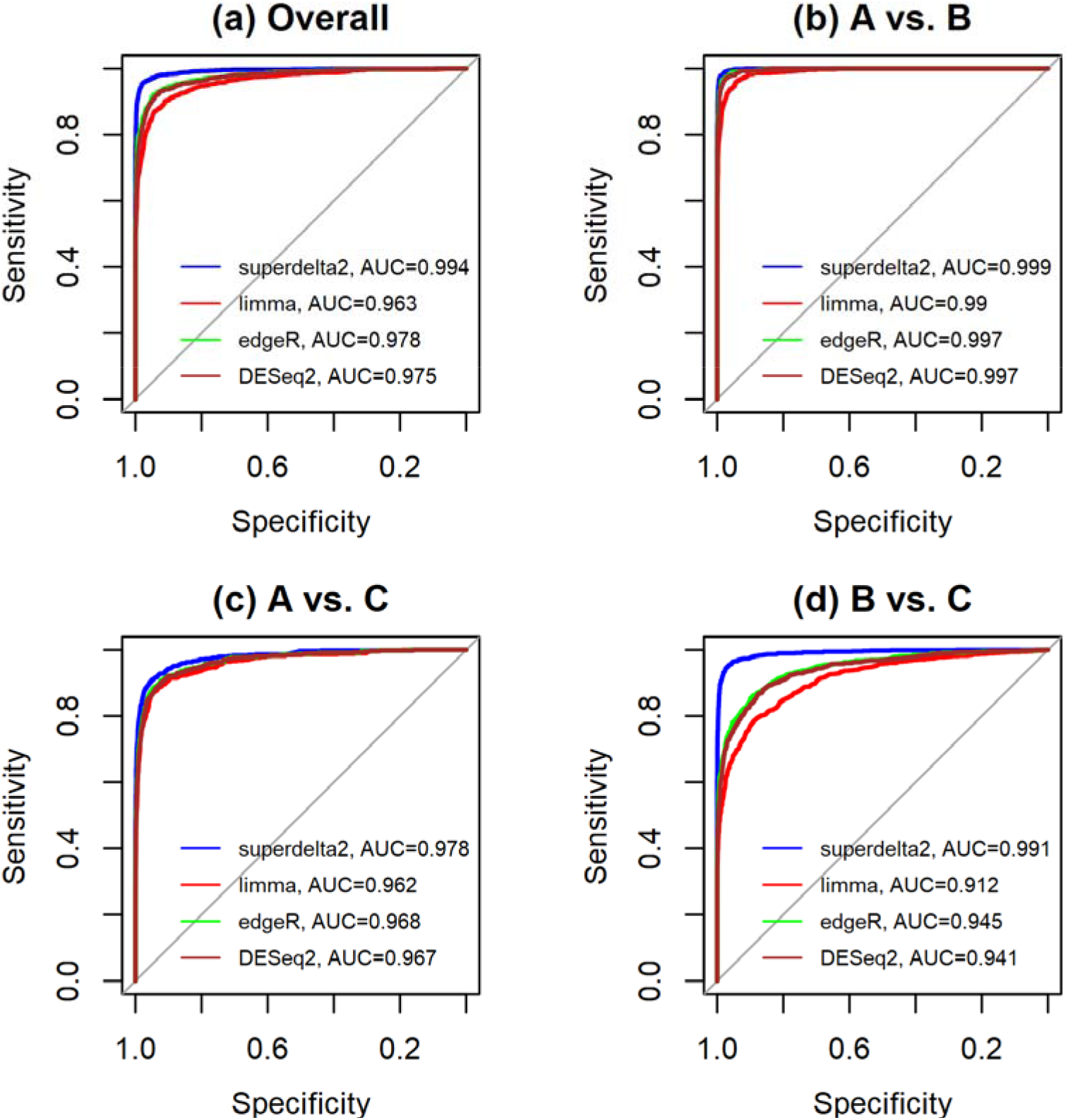
ROC curves of multi-group comparisons for simulation 1. (a) overall one-way ANOVA test; (b) Group A vs. Group B; (c) Group A vs. Group C; and (d) Group B vs. Group C.

We also recorded the computational time of each method. For a single run in simulation 1, `super-delta2` spent 13.0 seconds on average, which is slower than limma+voom (3.1 seconds), but is much faster than GLM-based algorithms DESeq2 (66.9 seconds) and edgeR (34.1 seconds).

### Simulation 2

In simulation 2, we assume that the read counts, *Y*_*i,gj*_, follows the following NBP model

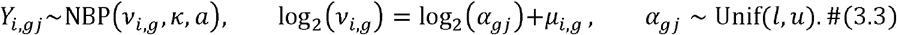

Unlike simulation 1, technical noise of the *j*th sample in group *g* (also denoted by *α*_*gj*_) is not used as a multiplicative constant (see Equation (3.2)), but as a latent factor that alters the conditional expectation of the read counts, 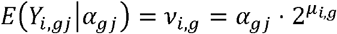. Unconditional group means are controlled by 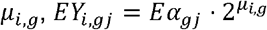.

In order to make simulations 1 and 2 comparable, we set *κ* = 0.06, *a* = 2.2, *l* = 12, and *u* = 30. Parameter *μ*_*i,g*_ is specified as follows:

1. Group A was set as the baseline, where *μ*_*i,A*_ = log_2_ 100, so *EY*_*i,gj*_ = 2100.
2. For Group B, genes 1-600 are up-regulated with *μ*_*i,B*_ = log_2_ 150, so *EY*_*i,gj*_ = 3150. Other genes have the same mean counts as group A (*v*_*i,B*_ = 100, *i* = 601,…5000).
3. For Group C, genes 401-1000 are down-regulated with *μ*_*i,C*_ = log_2_ 75, so *EY*_*i,gj*_ = 1575. Other genes have the same mean counts as group A.

The observed type I error rate and statistical power for each method are summarized in Table 2. Compared to simulation 1, the statistical power of simulation 2 decreases for all methods, but `super-delta2` is again the most powerful method in the overall multi-group comparison. More importantly, `super-delta2` still controlled type I error well. In fact, due to much tighter control of type I error, `super-delta2` tended to select fewer DEGs than other methods. For example, in one-way ANOVA test, on average `super-delta2` selected 1023 DEGs, which is fewer than all other methods (limma+voom: 1145, edgeR: 1144, DESeq2: 1214). See Appendix 4 in Supplementary Text for more details. This phenomenon has been observed in our real data analysis as well.

**Table 2:** Type I error rate, statistical power, and AUC values of multi-group comparisons at significance level *α* = 0.05 for various methods in **simulation 2**. The 3^rd^ column (Overall) records statistical performance of the one-way ANOVA test, the rest three columns record results from post-hoc pairwise group comparisons. All reported results are averaged over 100 repetitions. (·) represents the standard deviation of these 100 repetitions.

|  | Method | Overall | A vs. B | A vs. C | B vs. C |
| --- | --- | --- | --- | --- | --- |
| Type I error | `super-delta2` | **0.046**<br>(0.003) | **0.048**<br>(0.002) | **0.048**<br>(0.002) | **0.046**<br>(0.002) |
|  | `limma+voom` | 0.100<br>(0.005) | 0.081<br>(0.005) | 0.058<br>(0.003) | 0.122<br>(0.007) |
|  | `edgeR` | 0.083<br>(0.005) | 0.065<br>(0.004) | 0.055<br>(0.003) | 0.105<br>(0.006) |
|  | `DESeq2` | 0.098<br>(0.006) | 0.073<br>(0.004) | 0.064<br>(0.004) | 0.119<br>(0.007) |
| Statistical power | `super-delta2` | **0.839**<br>(0.009) | 0.909<br>(0.008) | 0.645<br>(0.010) | **0.817**<br>(0.009) |
|  | `limma+voom` | 0.745<br>(0.010) | 0.839<br>(0.009) | 0.595<br>(0.012) | 0.661<br>(0.011) |
|  | `edgeR` | 0.812<br>(0.009) | 0.916<br>(0.008) | 0.663<br>(0.011) | 0.744<br>(0.010) |
|  | `DESeq2` | 0.822<br>(0.008) | **0.922**<br>(0.007) | **0.677**<br>(0.010) | 0.753<br>(0.009) |
| AUC value | `super-delta2` | **0.966**<br>(0.003) | **0.984**<br>(0.001) | 0.912<br>(0.004) | **0.961**<br>(0.002) |
|  | limma+voom | 0.907<br>(0.004) | 0.957<br>(0.003) | 0.886<br>(0.006) | 0.851<br>(0.007) |
|  | edgeR | 0.942<br>(0.003) | 0.983<br>(0.001) | **0.916**<br>**(0.004)** | 0.903<br>(0.005) |
|  | DESeq2 | 0.940<br>(0.004) | 0.983<br>(0.001) | **0.916**<br>**(0.004)** | 0.899<br>(0.005) |

ROC curves in simulation 2 were provided in Figure 3. The AUC values are very similar to those in simulation 1, and `super-delta2` again has the highest overall AUC value.

**Figure 3:**
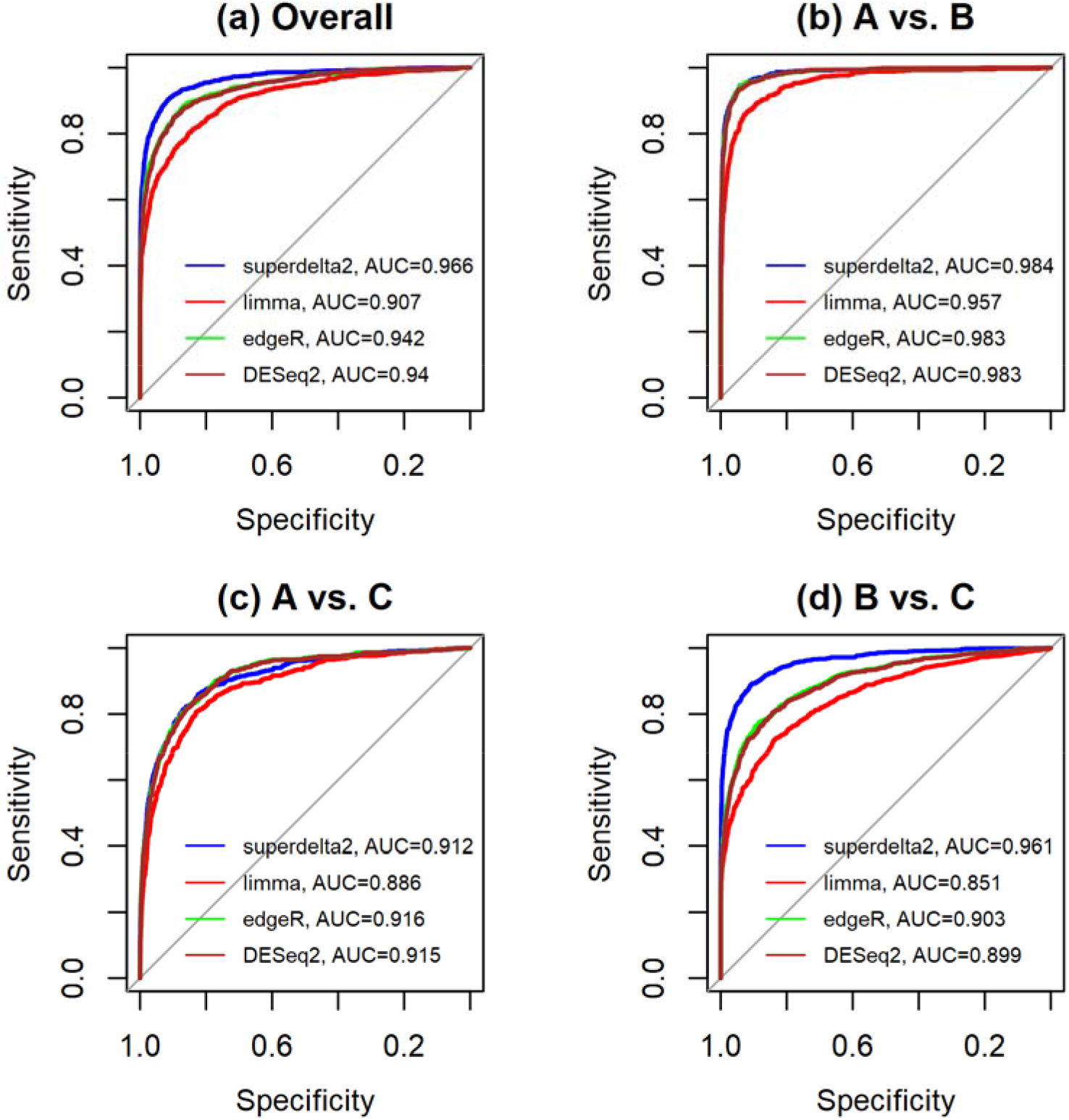
ROC curves of multi-group comparisons for simulation 2. (a) overall one-way ANOVA test; (b) Group A vs. Group B; (c) Group A vs. Group C; and (d) Group B vs. Group C.

### Simulation 3

In this simulation, we set *κ* = 0.6, *a* = 2, *l* = 10, and *u* = 20, to match those shape parameters estimated from the breast cancer data. Parameter *μ*_*i,g*_ is specified as follows:

1. Group A was set as the baseline, where *μ*_*i,A*_ = log_2_ 100, so *EY*_*i,gj*_ = 1500.
2. For Group B, genes 1-600 are up-regulated with *μ*_*i,B*_ = log_2_ 150, so *EY*_*i,gj*_ = 2250. Other genes have the same mean counts as group A (*v*_*i,B*_ = 100, *i* = 601,…5000).
3. For Group C, genes 401-1000 are down-regulated with *μ*_*i,C*_ = log_2_ 50, so *EY*_*i,gj*_ = 750. Other genes have the same mean counts as group A.

The results for simulation 3 were summarized in Table 3. Note that in this case, *a* = 2, so the negative binomial Poisson model is equivalent to negative binomial model, the statistical model used by DESeq2 and edgeR. This explains that in most cases, DESeq2 had the highest statistical power. However, DESeq2 had unacceptably high type I error in multi-group comparison and the pairwise comparison between groups B and C. `super-delta2` is the only method that controlled type I error tightly in all situations.

**Table 3:**
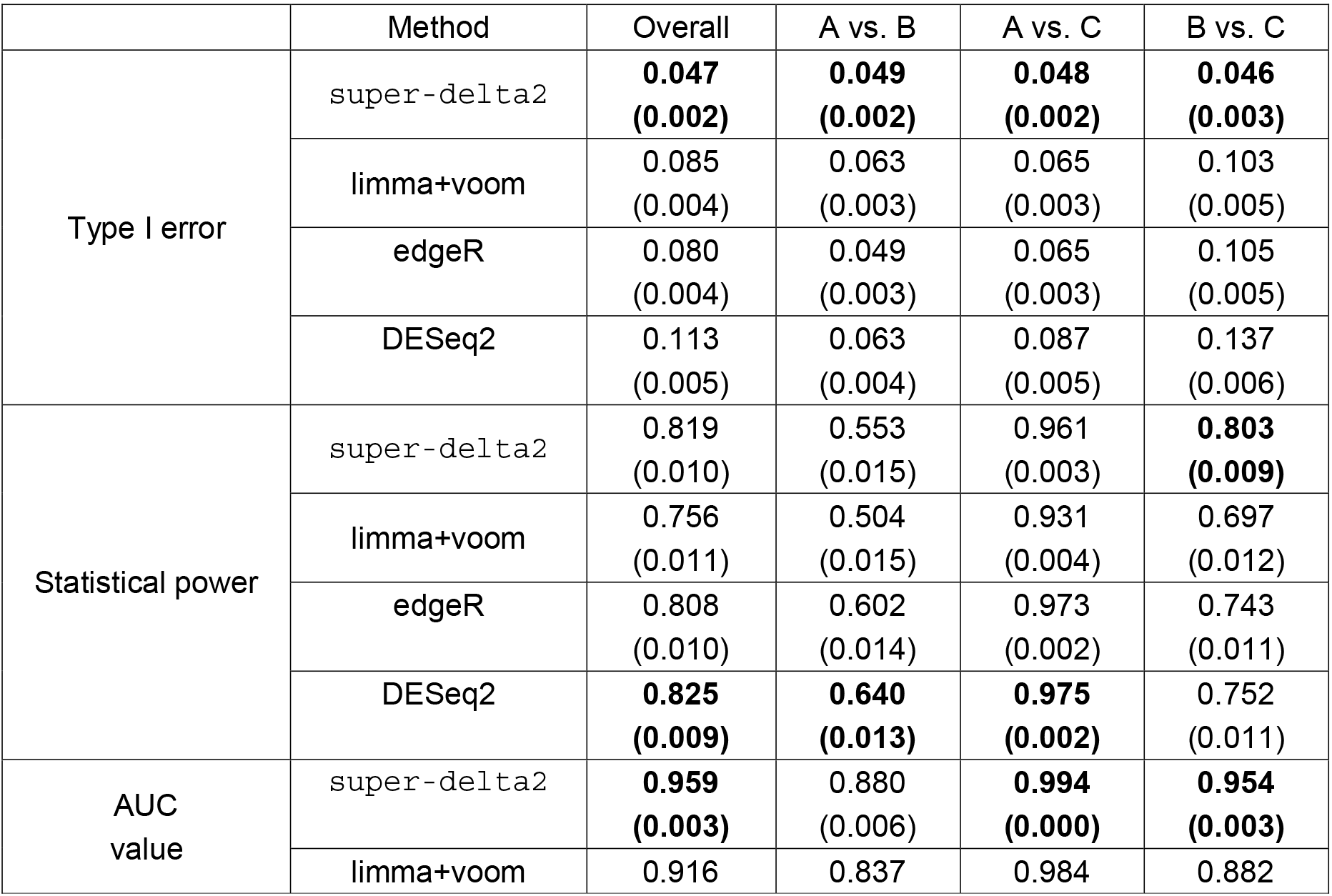
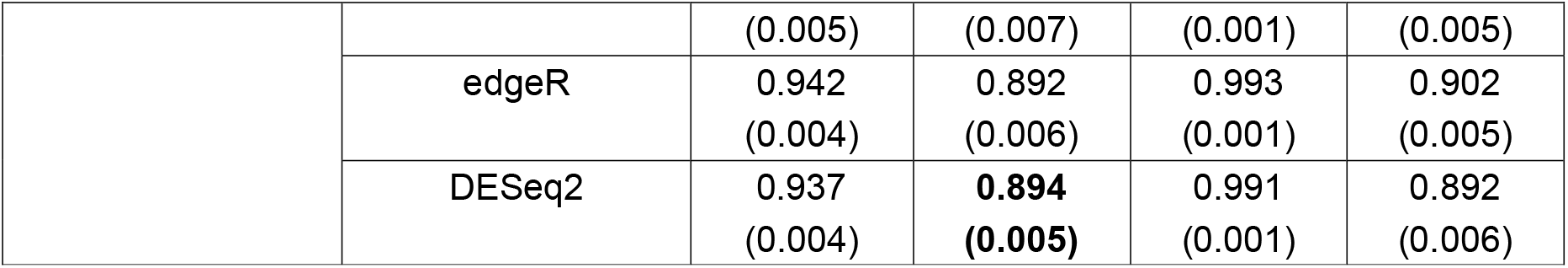
Type I error rate, statistical power, and AUC values of multi-group comparisons at significant level *α* = 0.05 for various methods in **simulation 3**. The 3^rd^ column (Overall) records statistical performance of the one-way ANOVA test, the rest three columns record results from post-hoc pairwise group comparisons. All reported results are averaged over 100 repetitions. (·) represents the standard deviation of these 100 repetitions.

ROC curves for simulation 3 are summarized in Figure 4. Again, `super-delta2` has the highest AUC value in overall comparisons.

**Figure 4:**
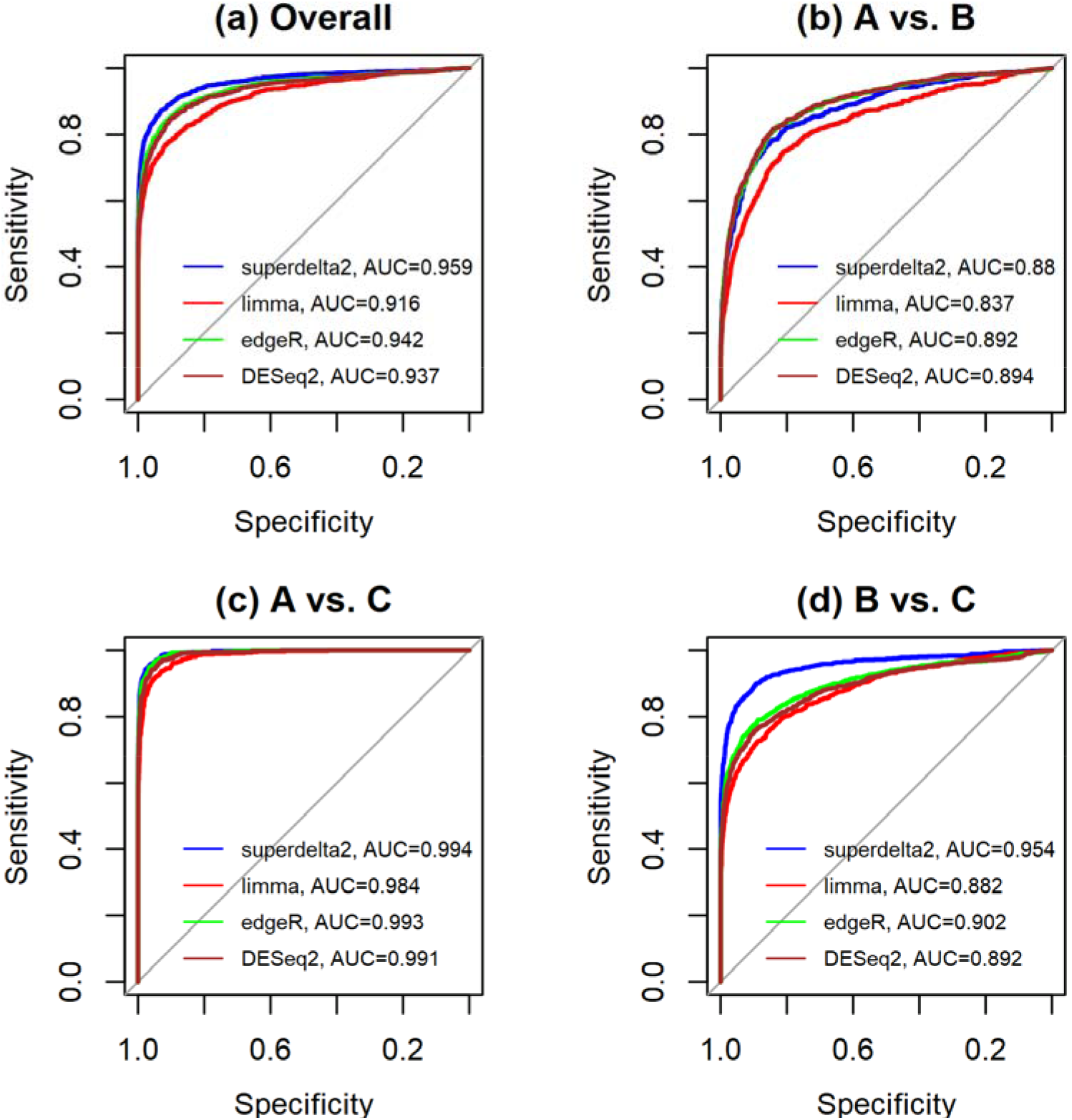
ROC curves of multi-group comparisons for simulation 3. (a) overall one-way ANOVA test, (b) Group A vs. Group B, (c) Group A vs. Group C, and (d) Group B vs. Group C.

We also tested the applicability of `super-delta2` for cases with limited observations by repeating the three simulations described above with only *n* = 10 samples in each group. By and large, the results are much the same: `super-delta2` was still the only method that controlled type I error at the nominal level, and it had the highest statistical power and AUC values in most cases. We even tried a more extreme case with only five samples in each group. Unfortunately, all four methods failed to control type I error, suggesting that we need *n* ≥ 10 samples per group in RNA-seq analysis to obtain reliable results. Technical details and results of these supplemental simulations are provided in Appendix 6.

Finally, we compared `super-delta2` with the original `super-delta` method for *post hoc* pairwise group tests in simulation 1. Both methods controlled type I error well, and `super-delta2` had slightly better statistical power and AUC values. More details are provided in Appendix 8.

## Real Data Analysis

### Differential Expression Analysis for the real data

The proposed `super-delta2` method, as well as three competitive methods (limma, DESeq2, and edgeR) were applied to a breast cancer (BRCA) dataset from The Cancer Genome Atlas (TCGA, https://cancergenome.nih.gov), which contains RNA-seq data of 1051 patients with tumor samples and their clinical information. Specifically, these patients were divided into three groups according to their pathologic stage (181 patients in stage I, 620 patients in stage II, and 250 patients in stage III). We applied CPM normalization (as implemented in edgeR) to filter genes at first. Genes with CPM value less than 1 on more than half of (526) patients were removed, the remaining 13,957 genes were used in subsequent DGEA. A gene is defined as differentially expressed if the unadjusted p-value in one-way ANOVA test is less than 0.05. Table 4 recorded the number of genes in significant gene lists from all four methods. We also recorded DEGs with BH-adjustment (adjusted p-value is less than 0.05) from these methods.

**Table 4:** Number of differential expressed genes detected by four methods (based on both unadjusted and BH-adjusted p-value) in one-way ANOVA test.

| Method | Number of DEGs | Number of DEGs with BH-adjustment |
| --- | --- | --- |
| DESeq2 | 4795 | 2816 |
| edgeR | 4679 | 2664 |
| Limma+voom | 3651 | 1201 |
| `super-delta2` | 3370 | 929 |

Based on our simulation analyses, we believe that the fact that `super-delta2` identified the smallest number of DEGs is likely due to its tighter control of type I error.

We applied pathway analyses to understand how these DEGs work together to dysregulate certain biological processes. Because the number of DEGs in the input can have strong effect on the pathway-level inference, we decide to use top 2,000 DEGs ranked by the p-values produced by all four DGEA methods as inputs for pathway analyses to remove this confounding effect. A Venn Diagram of these top 2,000 DEGs was created to show the number of overlapping genes between different methods (Figure 5). Although four methods produce prominently different list of DEGs, the similarities among them are also conspicuous. DESeq2 and edgeR, which both employ generalized linear model based on negative binomial distribution, have closer results (1768 genes in common), while limma and `super-delta2`, which both incorporate normality based linear models, tend to agree more (1784 genes in common).

**Figure 5:**
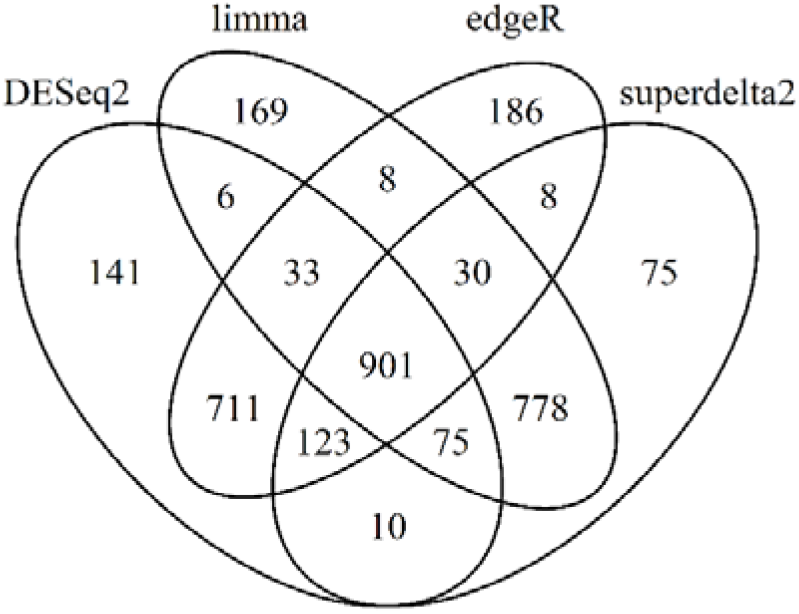
Venn Diagram of top 2,000 significant genes identified by four DGEA methods

### Gene Annotation and Gene Set Enrichment Analysis

We performed pathway analysis to those top 2,000 DEGs by Database for Annotation, Visualization and Integrated Discovery (DAVID version 6.8 [23]) with a focus on KEGG pathways [24]. Lists of significant KEGG pathways with Benjamini-Hochberg adjusted p-value less than 0.05 were shown in Table 5.

**Table 5:** Significant KEGG pathways selected by each DGEA method. (·) represents the number of top significant genes in the corresponding pathway.

| Method | Significant pathways |
| --- | --- |
| DESeq2 | hsa04110: Cell cycle (42) |
|  | hsa03030: DNA replication (15) |
|  | hsa03050: Proteasome (15) |
| edgeR | hsa04110: Cell cycle (43) |
|  | hsa03030: DNA replication (16) |
| limma+voom | hsa04110: Cell cycle (37) |
|  | hsa05200: Pathways in cancer (66) |
|  | hsa04114: Oocyte meiosis (27) |
|  | hsa04914: Progesterone-mediated oocyte maturation (21)<br>hsa04610: Complement and coagulation cascades (17) |
| super-delta2 | hsa04110: Cell cycle (40)<br>hsa03030: DNA replication (15)<br>hsa05200: Pathways in cancer (66)<br>hsa04114: Oocyte meiosis (27)<br>hsa03460: Fanconi anemia pathway (15)<br>hsa04914: Progesterone-mediated oocyte maturation (20)<br>hsa04151: PI3K-Akt signaling pathway (54) |

There are 7 significant KEGG pathways selected by `super-delta2`, the most among all four methods. Cell cycle (KEGG hsa04110), which was detected by all four methods, plays fundamental roles in controlling cell growth and death, therefore has close relation to the tumorigenesis, development, and metastasis of cancer. DNA replication (KEGG hsa03030), detected by DESeq2, edgeR, and `super-delta2`, is also a key biological process in tumor cell proliferation. Limma+voom and `super-delta2` identified more significant pathways, including Oocyte meiosis (has04114), Progesterone-mediated oocyte maturation (hsa04914), which are related to pathogenesis of breast cancer [25]; and Pathways in cancer (KEGG hsa05200), which consists of multiple known sub-pathways in cancer. Two pathways are selected only by `super-delta2`: Fanconi anemia pathway (KEGG hsa03460) and PI3K-Akt signaling pathway (KEGG hsa04151). One gene in Fanconi anemia pathway, FANCD1, is identical to a breast cancer susceptibility gene, BRCA2 [26]. Activation of PI3K-Akt signaling induces endocrine resistance in metastatic breast cancer, irrespective of the kind of endocrine agents administered [27]. Of note, other methods also identified some unique significant pathways: Proteasome (KEGG hsa03050), detected by DESeq2 and Complement and coagulation cascades (KEGG has 04610), detected by limma+voom. Overall, DESeq2 and edgeR found much less pathways related to cancer compared to `super-delta2` and limma+voom, and `super-delta2` identified the most number of pathways when 2000 DEGs were used for all methods. One explanation is that many genes selected by DESeq2 and edgeR may be false positives. This can also be reflected by the result of simulation study shown earlier, where DESeq2 and edgeR cannot control the type I error rate well.

### Rank Difference Analysis

We conduct a rank difference analysis to help understand why different methods made very different inference for certain genes. In this analysis, we focused on `super-delta2` and DESeq2 since they apply different models and DESeq2 is widely used in RNA-seq data analysis. In order to simplify this analysis, we focused on genes in Pathways in cancer (KEGG hsa05200). Genes were first ranked by their p-values from the smallest (most significant) to the largest (least significant) for both DESeq2 and `super-delta2`, respectively. We then calculated the difference of two ranks with larger absolute value of the rank difference indicating more disagreement between `super-delta2` and DESeq2. This analysis produced two types of genes: genes ranked high (significantly differentially expressed based on p-values) by DESeq2, but not `super-delta2`, and genes ranked high by `super-delta2`, but not DESeq2. Table 6 list top six genes with the largest absolute rank differences for each type, respectively.

**Table 6:** Six genes ranked high (significantly differentially expressed based on p-values) by DESeq2 (`super-delta2`), but not `super-delta2` (DESeq2), ordered by the absolute value of rank differences.

| Gene | Rank in DESeq2 | Rank in `super-delta2` | Rank difference | Gene | Rank in DESeq2 | Rank in `super-delta2` | Rank difference |
| --- | --- | --- | --- | --- | --- | --- | --- |
| WNT11 | 1777 | 13218 | -11441 | CBLC | 13794 | 1091 | 12703 |
| EGFR | 184 | 9625 | -9441 | HGF | 12888 | 786 | 12102 |
| PLCG2 | 859 | 8721 | -7862 | ARNT2 | 11683 | 1535 | 10148 |
| CDK6 | 928 | 8634 | -7706 | RUNX1T1 | 10622 | 1106 | 9516 |
| LAMA1 | 577 | 8004 | -7427 | FGF14 | 9761 | 1570 | 8191 |
| BMP2 | 1043 | 8261 | -7218 | FZD10 | 8263 | 199 | 8064 |

We selected three examples from Table 6 and drew boxplots of expression in each pathologic stage in Figure 6, in which the first two columns (CBLC, FZD10) were DEGs selected by `super-delta2` but not DESeq2; the last one (EGFR) was selected by DESeq2 but not `super-delta2`. We observe that CBLC, FZD10 have more differences among three stages than EGFR in terms of *median* expression levels. On the other hand, mean expressions of EGFR are very different among three stages, primarily due to outliers in stage II. This spurious group mean difference was picked up by DESeq2 but not `super-delta2`, because the majority of outliers were removed in the trimming process of `super-delta2`. which suggest that `super-delta2` is more robust to outliers. The boxplot of all genes in Table 6 are provided in appendix 5.

**Figure 6:**
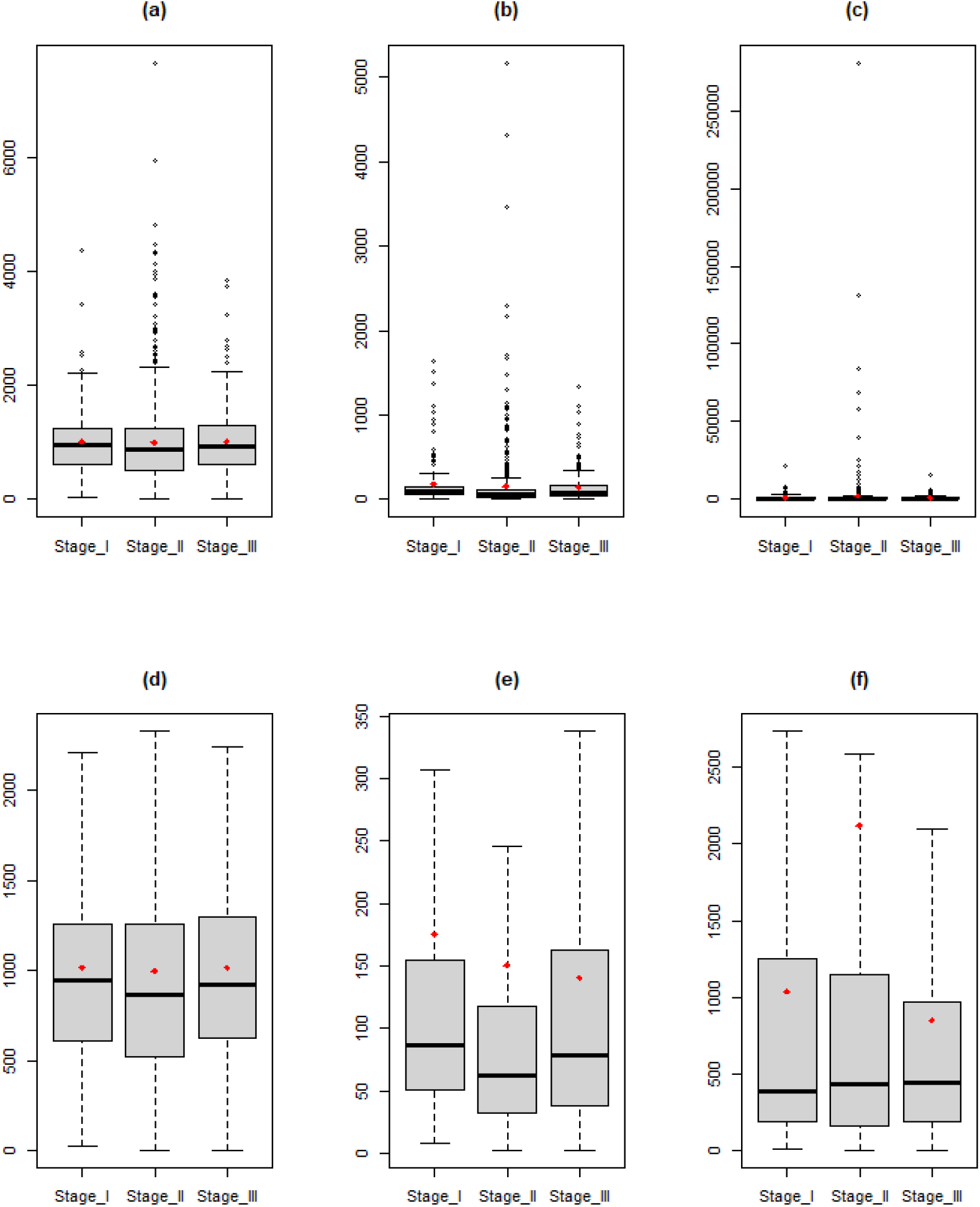
(a)-(c): Boxplot of RNA-seq data of CBLC, FZD10 and EGFR in three pathologic stages, respectively. (d)-(f) are the same plots as (a)-(c) without outliers. Red dots are stage-specific mean expression levels. The first two (CBLC and FZD10) are DEGs selected by `super-delta2` but not DESeq2. The last one (EGFR) was selected by DESeq2 but not `super-delta2`. Those two genes selected by `super-delta2` have more pronounced group differences than the one selected by DESeq2 in terms of median expression levels.

### Impact of Non-specific Filtering of Under-Expressed Genes

In this study, all four methods were conducted after removing genes with low mean expressions. The presence of these low-expression genes may decrease the sensitivity of DGEA. Thus, identifying and removing these genes prior to DGEA is a popular practice[13, 14, 22, 28, 29]. However, “under-expressed genes” is a relative concept, and different investigators may decide to use different thresholds in non-specific filtering, or even prefer to keep all genes for analysis. It is therefore important for a DGEA method to produce results that are consistent with or without filtering. With this motivation, we compared the top DEGs obtained with and without non-specific filtering by four methods. The results were summarized in Table 7.

**Table 7:** Number of common top DEGs (based on raw p-values) selected by each method in real data analysis with and without filtering out under-expressed genes. A total number of 20,531 genes were used in DGEA without filtering, and 13,957 genes were used in DGEA with filtering.

|  | super-delta2 | limma+voom | edgeR | DESeq2 |
| --- | --- | --- | --- | --- |
| Top 1000 | 772 | 795 | 38 | 372 |
| Top 2000 | 1507 | 1552 | 411 | 976 |
| Top 3000 | 2242 | 2295 | 968 | 1648 |

Based on Table 7, we see that both `super-delta2` and limma+voom produced consistent lists of DEGs with or without filtering. In comparison, the consistency of edgeR and DESeq2 was much lower, suggesting that these two GLM-based methods are more sensitive to different choices in pre-filtering and less reproducible. We also recorded the total number and the percentage of DEGs detected by all four methods with and without filtering in Table 8.

**Table 8:** Number of DEGs (raw p-value smaller than 0.05) selected with and without filtering by each method in real data analysis. (·) represents the percentage of DEGs detected by the corresponding setting.

|  | super-delta2 | limma+voom | edgeR | DESeq2 |
| --- | --- | --- | --- | --- |
| With filtering<br>(13957 genes) | 3370<br>(24.15%) | 3651<br>(26.16%) | 4679<br>(33.52%) | 4795<br>(34.36%) |
| Without filtering<br>(20531 genes) | 4502<br>(21.93%) | 4967<br>(24.19%) | 8089<br>(39.40%) | 7267<br>(35.40%) |

From Table 8, we find that the percentages of DEGs were higher after filtering for `super-delta2` and limma/voom, which is consistent with the common wisdom that on average, under-expressed genes contain less useful DEGs than other genes. However, edgeR and DESeq2 detected even more DEGs in under-expressed genes than fully-expressed genes, which is an indirect evidence of inflation of type I error for these two methods.

## Discussion

In summary, we proposed a differential gene expression analysis pipeline, named `super-delta2`, which is a generalization of its precedent `super-delta` [16] to one-way ANOVA multi-group comparisons. Compared with its predecessor, `super-delta2` has the following innovations.

First of all, we developed the modified ANOVA F-test to make the DGEA work with the robust multivariate extension of the CPM normalization. We also developed a Tukey style post-hoc t-tests specifically optimized for `super-delta2`, so that researchers can run the “overall test” and pairwise group comparisons in one step. Through extensive large sample analysis of `super-delta2` based on the Negative Binomial Poisson model, we demonstrated that the proposed method is asymptotically valid for RNA-seq data in addition to microarray data, which are approximately normally distributed and were the focus of the original `super-delta` paper. This theoretical prediction was validated in extensive simulation studies based on the NBP model, in which `super-delta2` is the only method that has tight control of type I error.

Secondly, we developed a bias-correction procedure after the spherical trimming, to compensate for the under-estimation of BGRSS (see Appendix 1 in Supplementary Text) and improve the statistical power of `super-delta2`. We believe this method could be adapted for other statistical problems, such as robust variance and covariance estimator based on trimming, in a future study.

Thirdly, the robustness of `super-delta2` is reflected in the fact that it produces consistent results with or without filtering under-expressed genes. This is due to the use of internal normalization and trimming in `super-delta2`. This is an additional advantage of our method because the type I error rate is not inflated when under-expressed genes are included in the DGEA due to biological reasons.

Last but not the least, we want to give plausible explanations to the observation that all three other methods failed to control type I error at the nominal level in simulation studies, especially when the statistical power is high (simulation 1). All three methods have internal data transformations that can be considered as variants of traditional variance stabilizing and normalization procedures. These methods reduce the variability in the data by borrowing information from both differentially expressed and non-differential expressed genes, as well as genes with outliers. Such practice inevitably introduces some artificial bias when the between group difference of up-regulated and down-regulated genes is complex, especially when the structure is unbalanced [7–9]. On the other hand, the trimmed multivariate normalization procedure in `super-delta2` effectively removes these detrimental effects thus has the best control of type I error. We must admit that all trimming methods remove a small proportion of data thus may reduce statistical power slightly. By design, trimming in `super-delta2` removes genes, not samples. Since there are many more genes than samples in a typical genomics study, this strategy preserves the statistical power well, as shown in our simulation studies. Furthermore, pathway analyses of the real data showed that identifying a smaller but more accurate set of DEGs can often help the biologist identify richer and more relevant information. Hence, we think it is always a good idea to use trimming in practice.

A minor point is that, unlike `super-delta2` that has closed-form formula for conducting hypothesis tests, edgeR and DESeq2 depend on iterative nonlinear optimization procedures to fit GLMs first. Occasionally, these numerical algorithms fail to converge to global maxima, which may affect the subsequent hypothesis tests.

In the future, we believe it would be fruitful to generalize `super-delta2` so that it works with multiple regressions, possibly with both fixed and random effects. This requires us to design multivariate normalization (the *δ*-step) that works for multiple regression, a trimming based on Mahalanobis distance instead of Euclidean distance of the equivalence of “*R*_*ik,g*_” that was used in estimating BGRSS in `super-delta2`, and possibly a bias-correction procedure for this trimming.

## Author Contributions

XQ and JZ conceptualized and designed the study. ZC and YL performed data analyses and interpreted results. All four authors wrote the manuscript and approved the final version.

## Availability of data and material

Our method is implemented in a R-package, “`superdelta2`”, freely available at: https://github.com/fhlsjs/superdelta2

## Acknowledgements

This work was supported by the National Institute of General Medical Sciences of the National Institute of Health under award number R01GM126558 for JZ and the University of Rochester CTSA award number UL1 TR002001 from the National Center for Advancing Translational Sciences of the National Institutes of Health for XQ. The funders had no role in study design, data collection and analysis, decision to publish, or preparation of the manuscript.

## References

1. Yang YH, Dudoit S, Luu P, Lin DM, Peng V, Ngai J, Speed TP: Normalization for cDNA microarray data: a robust composite method addressing single and multiple slide systematic variation. Nucleic Acids Res 2002, 30(4):e15.

2. Parrish RS, Spencer III HJ: Effect of normalization on significance testing for oligonucleotide microarrays. Journal of biopharmaceutical statistics 2004, 14(3):575–589.

3. Bolstad BM, Irizarry RA, Astrand M, Speed TP: A comparison of normalization methods for high density oligonucleotide array data based on variance and bias. Bioinformatics 2003, 19(2):185–193.

4. Robinson MD, Oshlack A: A scaling normalization method for differential expression analysis of RNA-seq data. Genome Biol 2010, 11(3):R25.

5. Roberts A, Trapnell C, Donaghey J, Rinn JL, Pachter L: Improving RNA-Seq expression estimates by correcting for fragment bias. Genome Biol 2011, 12(3):R22.

6. Hansen KD, Irizarry RA, Wu Z: Removing technical variability in RNA-seq data using conditional quantile normalization. Biostatistics 2012.

7. Qiu X, Hu R, Wu Z: Evaluation of Bias-variance Trade-off for Post-summarizing Normalization Procedures in Large-Scale Genomic Studies. PloS One 2014, 9(6):e99380.

8. Qiu X, Wu H, Hu R: The impact of quantile and rank normalization procedures on the testing power of gene differential expression analysis. BMC Bioinformatics 2013, 14:124.

9. Liu Y, Zhang J, Qiu X: Super-delta: a new differential gene expression analysis procedure with robust data normalization. BMC Bioinformatics 2017, 18(1):582.

10. Di Y, Schafer DW, Cumbie JS, Chang JH: The NBP negative binomial model for assessing differential gene expression from RNA-Seq. Statistical applications in genetics and molecular biology 2011, 10(1).

11. Smyth GK: Linear models and empirical bayes methods for assessing differential expression in microarray experiments. Stat Appl Genet Mol Biol 2004, 3:Article3.

12. Law CW, Chen Y, Shi W, Smyth GK: voom: Precision weights unlock linear model analysis tools for RNA-seq read counts. Genome Biol 2014, 15(2):R29.

13. Robinson MD, McCarthy DJ, Smyth GK: edgeR: a Bioconductor package for differential expression analysis of digital gene expression data. Bioinformatics 2010, 26(1):139–140.

14. Love M, Anders S, Huber W: Differential analysis of count data–the DESeq2 package. Genome Biol 2014, 15(550):10–1186.

15. Tsodikov A, Szabo A, Jones D: Adjustments and measures of differential expression for microarray data. Bioinformatics 2002, 18(2):251–260.

16. Ni TT, Lemon WJ, Shyr Y, Zhong TP: Use of normalization methods for analysis of microarrays containing a high degree of gene effects. BMC bioinformatics 2008, 9(1):505.

17. Qin L-X, Satagopan JM: Normalization method for transcriptional studies of heterogeneous samples-simultaneous array normalization and identification of equivalent expression. Statistical applications in genetics and molecular biology 2009, 8(1):1–23.

18. Ogunnaike BA, Gelmi CA, Edwards JS: A probabilistic framework for microarray data analysis: Fundamental probability models and statistical inference. Journal of theoretical biology 2010, 264(2):211–222.

19. Anders S, Huber W: Differential expression analysis for sequence count data. Genome Biol 2010, 11(10):R106.

20. Marioni JC, Mason CE, Mane SM, Stephens M, Gilad Y: RNA-seq: an assessment of technical reproducibility and comparison with gene expression arrays. Genome Res 2008, 18(9):1509–1517.

21. Rapaport F, Khanin R, Liang Y, Pirun M, Krek A, Zumbo P, Mason CE, Socci ND, Betel D: Comprehensive evaluation of differential gene expression analysis methods for RNA-seq data. Genome Biol 2013, 14(9):R95.

22. Ritchie ME, Phipson B, Wu D, Hu Y, Law CW, Shi W, Smyth GK: limma powers differential expression analyses for RNA-sequencing and microarray studies. Nucleic acids research 2015, 43(7):e47–e47.

23. Sherman BT, Lempicki RAJNp: Systematic and integrative analysis of large gene lists using DAVID bioinformatics resources. 2009, 4(1):44.

24. Kanehisa M, Goto S: KEGG: kyoto encyclopedia of genes and genomes. Nucleic Acids Res 2000, 28(1):27–30.

25. Wu D, Han B, Guo L, Fan ZJJoO, Gynaecology: Molecular mechanisms associated with breast cancer based on integrated gene expression profiling by bioinformatics analysis. 2016, 36(5):615–621.

26. D’Andrea AD: Susceptibility pathways in Fanconi’s anemia and breast cancer. The New England journal of medicine 2010, 362(20):1909–1919.

27. Tokunaga E, Kimura Y, Mashino K, Oki E, Kataoka A, Ohno S, Morita M, Kakeji Y, Baba H, Maehara YJBc: Activation of PI3K/Akt signaling and hormone resistance in breast cancer. 2006, 13(2):137–144.

28. Love MI, Huber W, Anders S: Moderated estimation of fold change and dispersion for RNA-seq data with DESeq2. Genome Biol 2014, 15(12):550.

29. Bourgon R, Gentleman R, Huber W: Independent filtering increases detection power for high-throughput experiments. Proc Natl Acad Sci U S A 2010, 107(21):9546–9551.

