## Supplemental Text for "Super-delta2: An Enhanced Differential Expression Analysis Procedure for Multi-Group Comparisons of RNA-seq Data"

### Appendix 1: Bias Correction for Spherical Trimming

As a reminder, the BGRSS used in `super-delta2` F-test is estimated from between-group residuals computed from the deltas by Equation (2.9) in the main text:

$$R_{ik,g} := \sqrt{N_g}(\bar{\delta}_{ik,g\cdot} - \bar{\bar{\delta}}_{ik,\cdot\cdot}) = \sqrt{N_g}\left(\mu_{i,g} - \mu_{k,g} + \bar{\epsilon}_{i,g\cdot} - \bar{\epsilon}_{k,g\cdot} - \left(\bar{\mu}_{i,\cdot} - \bar{\mu}_{k,\cdot} + \bar{\bar{\epsilon}}_{i,\cdot\cdot} - \bar{\bar{\epsilon}}_{k,\cdot\cdot}\right)\right)$$
$$= \underbrace{\sqrt{N_g}\left(\mu_{i,g} + \bar{\epsilon}_{i,g\cdot} - \left(\bar{\mu}_{i,\cdot} + \bar{\bar{\epsilon}}_{i,\cdot\cdot}\right)\right)}_{R^*_{i,g}\text{: the oracle part}} - \underbrace{\sqrt{N_g}\left(\mu_{k,g} - \bar{\mu}_{k,\cdot}\right)}_{\text{possible bias}} + \underbrace{\sqrt{N_g}\left(\bar{\epsilon}_{k,g\cdot} - \bar{\bar{\epsilon}}_{k,\cdot\cdot}\right)}_{\text{additional variation}}.$$

Let us denote $R_{ik,g}$ collectively as a vector $\boldsymbol{R}_{ik} := \left(R_{ik,1}, \dots R_{ik,G}\right) \in \mathbb{R}^G$, and the second term in the above equation by $\boldsymbol{B}_k := \sqrt{N_g}\left(\mu_{k,1} - \bar{\mu}_{k,\cdot}, \dots \mu_{k,G} - \bar{\mu}_{k,\cdot}\right)' \in \mathbb{R}^G$. When $k \in \mathcal{S}^0$, $\boldsymbol{B}_k = 0_G$ therefore $R_{ik,g}$ is an unbiased predictor of $R^*_{i,g}$ (the first term in the above Equation). When $k \in \mathcal{S}^1$, we know that all group means are not equal so $\boldsymbol{B}_k \neq 0_G$. This bias may deteriorate the subsequent tests, therefore we decided to use trimming technique to remove those $\boldsymbol{R}_{ik}$ with extremely large squared Euclidean length $L_{ik} := \sum_{g=1}^G R^2_{ik,g}$. Specifically, For a pre-specified $q \in (0,1)$, let $L_{i,(m_q)}$, $m_q := [(m-1)(1-q)]$, be the $(1-q)$th empirical quantile of $L_{i\cdot}$. We define an index set $S_{i,q} := \left\{k = 1,2,\dots,m;\ k \neq i;\ L_{ik} \leq L_{i,(m_q)}\right\}$, and define the trimmed estimator of BGRSS as follows

$$R^{\text{trim}}_{i,g} := \frac{1}{|S_{i,q}|}\sum_{k\in S_q} R_{ik,g}\,,\quad \widehat{\text{BGRSS}}_i := \sum_{g=1}^G \left(R^{\text{trim}}_{i,g}\right)^2 = \left|R^{\text{trim}}_{i,g}\right|^2.$$

However, this method will inevitably under-estimate BGRSS because after trimming, $\boldsymbol{R}_{ik}$ (for $k \in S_{i,q}$) follows a *spherically trimmed multivariate normal* (STMVN) distribution, and the sample mean of the remaining observations, $\frac{1}{|S_{i,q}|}\sum_{k\in S_q} R_{ik,g}$, is no longer an unbiased estimator of the conditional expectation $E\left(R_{ik,g}\middle|\bar{\epsilon}_{i,g\cdot}\right) = \sqrt{N_g}\left(\mu_{i,g} + \bar{\epsilon}_{i,g\cdot} - \left(\bar{\mu}_{i,\cdot} + \bar{\bar{\epsilon}}_{i,\cdot\cdot}\right)\right)$. In fact, in most cases, $\widehat{\text{BGRSS}}_i = \left|R^{\text{trim}}_{i,g}\right|^2$ tend to *under-estimate* the oracle BGRSS (the squared Euclidean length of $E\left(R_{ik,g}\middle|\bar{\epsilon}_{i,g\cdot}\right)$), because trimming not only removes outliers, but also non-outliers with large Euclidean distances.

We aim to develop a bias-correction procedure based on the moment method for the STMVN in this section. We will start with a special case of STMVN first. Let $\mathbf{e}_1 := (1,0,\dots,0)'$ and $\mathbf{X} = (X_1, X_2, \dots, X_K)' \sim N(\mu \cdot \mathbf{e}_1,\ I_{K\times K})$, for $\mu \geq 0$, be a $K$-dimensional multivariate normal distribution. Note that, by construction, only the first coordinate of $\mathbf{X}$ has a nonzero mean. Denote by $D_R := \{\mathbf{x} \in \mathbb{R}^K : \|\ \mathbf{x}\ \| \leq R\}$ a ball with radius $R$ centered at the origin, and we consider a spherical trimming of $\mathbf{X}$ with acceptance domain $D_R$. The expectation of the remaining observations, which can be considered as realizations sampled from an STMVN, equals the conditional expectation $E(\mathbf{X}|D_R)$ that can be represented as follows.

$$E(\mathbf{X}|D_R) = S_R(\mu)\cdot \mathbf{e}_1, \qquad E(X_k|D_R) = \begin{cases} S_R(\mu), & k = 1, \\ 0, & k \neq 1. \end{cases} \tag{A1.1}$$

Here function $S_R(\mu) := E(X_1|\mathbf{X} \in D_R)$ is defined by

$$S_R(\mu) := \frac{\int_{-R}^R x \cdot F_{\chi^2_{K-1}}(R^2 - x^2)\cdot \phi(x-\mu)\mathrm{d}x}{\int_{-R}^R F_{\chi^2_{K-1}}(R^2 - x^2)\cdot \phi(x-\mu)\mathrm{d}x} = \frac{\int_{-R}^R x e^{-(x-\mu)^2/2}\cdot \gamma\left(\frac{K-1}{2}, R^2 - x^2\right)\mathrm{d}x}{\int_{-R}^R e^{-(x-\mu)^2/2}\cdot \gamma\left(\frac{K-1}{2}, R^2 - x^2\right)\mathrm{d}x}. \tag{A1.2}$$

where $F_{\chi^2_{K-1}}(\cdot)$ is the distribution function of a $\chi^2$-distribution with $K-1$ degrees of freedom and $\gamma(\cdot,\cdot)$ is the lower incomplete gamma function. It can be easily seen from the above representation that $\frac{S_R(\mu)}{\mu} \in (0,1)$, and it can be considered as an attenuation factor of the expectation due of trimming.

As a remark, while $S_R(\mu)$ does not have a closed-form, it can be easily computed by a suitable numerical integration method.

Next, let us consider a more general case in which $\mathbf{X} \sim N(\boldsymbol{\mu},\ \sigma^2 I_{K\times K})$ for an arbitrary $\boldsymbol{\mu} \in \mathbb{R}^{\boldsymbol{K}}$ and $\sigma \in \mathbb{R}^+$. We will use a suitable rotation and dilation to map it to the special case we mentioned earlier. Specifically, Let $T$ be a rotation such that $T\boldsymbol{\mu} = \parallel \boldsymbol{\mu} \parallel \mathbf{e}_1$ and $T^{-1}\mathbf{e}_1 = \frac{\boldsymbol{\mu}}{\parallel\boldsymbol{\mu}\parallel}$. Let $\mathbf{Y} := \sigma^{-1}T\mathbf{X}$, it follows distribution $N((\parallel \boldsymbol{\mu} \parallel /\sigma, 0, \dots, 0)',\ I_{K\times K})$, and $\sigma^{-1}TD_R = D_{R/\sigma}$, which is a special case mentioned earlier. Based on this observation, we come to the following main conclusion:

**Theorem 1.1**: Let $\mathbf{X} = (X_1, X_2, \dots, X_K)' \sim N(\boldsymbol{\mu},\ \sigma^2 I_{K\times K})$ be a $K$-dimensional multivariate normal distribution. We have

$$E(\mathbf{X}|D_R) = \sigma \cdot S_{R/\sigma}(\parallel \boldsymbol{\mu} \parallel/\sigma) \cdot \frac{\boldsymbol{\mu}}{|\boldsymbol{\mu}|}, \qquad \frac{|E(\mathbf{X}|D_R)|}{\sigma} = S_{R/\sigma}(|\boldsymbol{\mu}|/\sigma).$$

**Proof**:

Using Equations (A3.1) and (A3.2), we have

$$E(\sigma^{-1}T\mathbf{X}|D_R) := E(\mathbf{Y}|D_R) = S_{R/\sigma}(\parallel \boldsymbol{\mu} \parallel/\sigma) \cdot \mathbf{e}_1,$$

which implies

$$E(\mathbf{X}|D_R) = \sigma T^{-1} E(\sigma^{-1}T\mathbf{X}|D_R) = \sigma \cdot S_{R/\sigma}(|\boldsymbol{\mu}|/\sigma) \cdot \frac{\boldsymbol{\mu}}{|\boldsymbol{\mu}|}.$$

Theorem 3.1 shows that as a vector in $\mathbb{R}^K$, $E(\mathbf{X}|D_R)$ shares the same _direction_ as $\boldsymbol{\mu}$ but with a different length: $|E(\mathbf{X}|D_R)| = \sigma \cdot S_{R/\sigma}(|\boldsymbol{\mu}|/\sigma)$ instead of $|\boldsymbol{\mu}|$. Conversely, if we know $|E(\mathbf{X}|D_R)|$, $R$, and $\sigma$, $|\boldsymbol{\mu}|$ can be obtained by the following equation

$$|\boldsymbol{\mu}| = \sigma \cdot S^{-1}_{R/\sigma}(|E(\mathbf{X}|D_R)|/\sigma) \ . \qquad \text{(A1.3)}$$

We implemented the inverse function $S^{-1}_{R/\sigma}(\cdot)$ by a numerical algorithm based on the Newton-Raphson method in our R package. Specifically, we first obtain trimmed group means, $R^{\text{trim}}_{i,g}$ and its Euclidean norm $|R^{\text{trim}}_{i,g}|$. We then replace $|E(\mathbf{X}|D_R)|$ in Equation (A3.3) by $|R^{\text{trim}}_{i,g}|$, and use the numerical inverse algorithm to find an estimate of $|\boldsymbol{\mu}|$; the square of this estimate, $\widehat{|\boldsymbol{\mu}|}^2$, is considered as the _bias-corrected_ estimation of $\text{BGRSS}_i$.

For convenience, below we derive the derivative of $S_R(\mu)$, which is needed in the Newton-Raphson method.

**Theorem 1.2**: Let us denote the top and bottom of $S_R(\mu)$ defined in Equation (A3.2) as

$$T_R(\mu) := \int_{-R}^{R} x\, e^{-(x-\mu)^2/2} \cdot \gamma\left(\frac{K-1}{2}, R^2 - x^2\right) \mathrm{d}x, \quad B_R(\mu) := \int_{-R}^{R} e^{-(x-\mu)^2/2} \cdot \gamma\left(\frac{K-1}{2}, R^2 - x^2\right) \mathrm{d}x.$$

In addition, we define

$$H_R(\mu) := \int_{-R}^{R} x^2\, e^{-(x-\mu)^2/2} \cdot \gamma\left(\frac{K-1}{2}, R^2 - x^2\right) \mathrm{d}x.$$

The derivative of $S_R(\mu)$ evaluated at $\mu$ can be represented as

$$\frac{\mathrm{d}S_R(\mu)}{\mathrm{d}\mu} = \frac{H_R(\mu)B_R(\mu) - T_R^2(\mu)}{B_R^2(\mu)}.$$

**Proof:**

The derivatives of $T_R(\mu)$ and $B_R(\mu)$ are

$$
\begin{aligned}
\frac{\mathrm{d}T_R(\mu)}{\mathrm{d}\mu} &= \int_{-R}^{R} x\,(x-\mu)e^{-(x-\mu)^2/2}\cdot\gamma\left(\tfrac{K-1}{2},\,R^2-x^2\right)\mathrm{d}x \\
&= \int_{-R}^{R} x^2\,e^{-(x-\mu)^2/2}\cdot\gamma\left(\tfrac{K-1}{2},\,R^2-x^2\right)\mathrm{d}x-\mu T_R(\mu) \\
&= H_R(\mu)-\mu T_R(\mu). \\
\frac{\mathrm{d}B_R(\mu)}{\mathrm{d}\mu} &= \int_{-R}^{R}(x-\mu)e^{-(x-\mu)^2/2}\cdot\gamma\left(\tfrac{K-1}{2},\,R^2-x^2\right)\mathrm{d}x \\
&= T_R(\mu)-\mu B_R(\mu).
\end{aligned}
$$

Therefore, the derivative of $S_R(\mu)$ is

$$
\begin{aligned}
\frac{\mathrm{d}S_R(\mu)}{\mathrm{d}\mu} &= \frac{T_R{}'(\mu)B_R(\mu)-T_R(\mu)B_R{}'(\mu)}{B_R(\mu)^2} \\
&= \frac{(H_R(\mu)-\mu T_R(\mu))B_R(\mu)-T_R(\mu)(T_R(\mu)-\mu B_R(\mu))}{B_R^2(\mu)} \\
&= \frac{H_R(\mu)B_R(\mu)-T_R^2(\mu)}{B_R^2(\mu)}.
\end{aligned}
$$

Finally, we created Figure S1 to illustrate the effect of this bias-correction for spherically trimmed bivariate normal random vector.

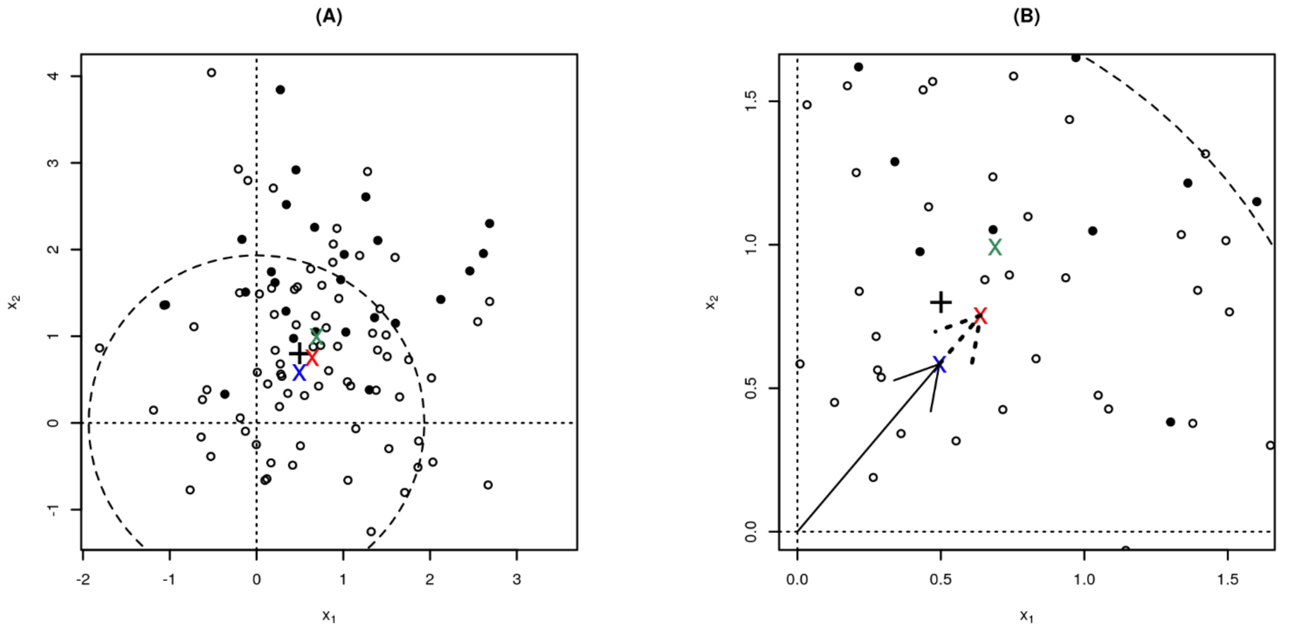


**Figure S1**: A graphical illustration of the bias-correction procedure. $n = 75$ circles are $X_i$ sampled from a 2D normal random vector with mean vector $\mu = (0.5, 0.8)$ and covariance matrix $\Sigma = I$. $n = 25$ solid dots represent outliers sampled from a 2D normal vector with $\mu^{\text{out}} = (0.9,2.0)$ and $\Sigma = I$. The circle represents a spherical trimming that filters out 30% of data points with the largest Euclidean length (greater than the radius of the circle, $R = 1.934$). Black "+" represents the true mean vector, "x" are the estimations of $\mu$: (a) the green one represents sample mean, $\bar{X} = (0.690, 0.992)$, with Euclidean length $|\bar{X}| = 1.209$, which is an over-estimate of $|\mu| = 0.943$ due to the presence of outliers, (b) the blue one represents trimmed mean of $X$, $\bar{X}^{\text{trim}} = c(0.495,0.584)$, with length $\left|\bar{X}^{\text{trim}}\right| = 0.765$, which under-estimates $|\mu|$ because the sample mean of the remaining observations is not an unbiased estimator of the location parameter of a spherically trimmed normal vector, and (c) the red one represents bias-corrected trimmed mean, $\bar{X}^{\text{corrected}} = c(0.639,0.754)$, with length $\left|\bar{X}^{\text{corrected}}\right| = 0.988$, which is the most accurate estimator of $|\mu|$ among three estimators. (A) shows all observations, and (B) is a zoomed in version that highlights the differences between the three estimators. Note that the bias-corrected estimator, $\bar{X}^{\text{corrected}}$, is a scaled version of $\bar{X}^{\text{trim}}$, so they have the same directions. This scaling transformation is represented by the two arrows in (B).

### Appendix 2: Large Sample Properties of `super-delta2` under Negative Binomial Poisson Model

First, let us recall that a random variable with negative binomial (NB) distribution has probability density function defined for integers $x = 0, 1, 2, \ldots$

$$\Pr(X = x|\nu, \gamma) = \frac{\Gamma(\gamma+x)}{\Gamma(\gamma)\Gamma(1+x)}\left(\frac{\nu}{\nu+\gamma}\right)^x\left(\frac{\gamma}{\nu+\gamma}\right)^\gamma. \tag{A2.1}$$

Here $\nu$ is location parameter and $\gamma$ is shape parameter. The mean and variance of the above NB distribution are

$$EX = \nu, \quad \mathrm{var}(X) = \nu + \frac{\nu^2}{\gamma}. \tag{A2.2}$$

According to [1], NBP model can be considered as an extension of negative binomial (NB) model. Three parameters are used in NBP model: the same location parameter $\nu$, as well as $\kappa$ and $a$ that link $\nu$ with the shape parameter $\gamma$ in the NB model with the following formula

$$\gamma := \kappa^{-1}\nu^{2-a}. \tag{A2.3}$$

the mean and variance of $X \sim \mathrm{NBP}(\nu, \kappa, a)$ are

$$EX = \nu, \quad \mathrm{var}(X) = \nu + \frac{\nu^2}{\gamma} = \nu + \kappa \cdot \nu^a. \tag{A2.4}$$

By comparing Equation (A2.4) with (A2.2), it is easy to see that the NB distribution is a *special case* of the NBP model with $\kappa = 1/\gamma$ and $a = 2$.

In practice, we also need to consider the impact of sample-specific technical noise to the observed gene expressions. We tried two approaches in this study. The following model is used by Simulation 1 (see Equation (3.2) in the main text):

$$X_{i,gj} \sim \mathrm{NBP}(\nu_{i,g}, \kappa, a), \; Y_{i,gj} = [\alpha_{gj} \cdot X_{i,gj}], \; \alpha_{gj} \sim F_\alpha. \tag{A2.5}$$

Here $[x]$ stands for the integer rounding function. Another model is used in Simulations 2 and 3 (see Equation (3.3) in the main text):

$$Y_{i,gj} \sim \mathrm{NBP}(\nu_{i,g}, \kappa_i, a_i), \; \log_2(\nu_{i,g}) = \log_2 \alpha_j + \mu_{i,g}, \; \alpha_{gj} \sim F_\alpha. \tag{A2.6}$$

In both models, $\alpha_j \sim F_\alpha$ is a latent random variable that represents technical noise. Due to the fact that the observed read counts are non-negative, $\alpha_j$ must be a non-negative random variable. In our simulations, $F_\alpha$ is set to be a uniform distribution. In this section, we would like to show that $F_\alpha$ can be a flexible distribution that satisfies relatively mild mathematical conditions, because we will prove that the effect of $\alpha_j$ is essentially removed by the $\delta$-step in `super-delta2`.

Let

$$W_{i,gj} := \begin{cases} \log_2(Y_{i,gj}), & Y_{i,gj} \geq 1 \\ 0, & Y_{i,gj} = 0 \end{cases}, \qquad \bar{W}_{i,g\cdot} := \frac{1}{N_g}\sum_{j=1}^{N_g} W_{i,gj};$$

$$\delta_{ik,gj} := W_{i,gj} - W_{k,gj}, \quad \bar{\delta}_{ik,g\cdot} := \frac{1}{N_g}\sum_{j=1}^{N_g}(W_{i,gj} - W_{k,gj}). \tag{A2.7}$$

For Model (A2.5), $W_{i,gj} \approx \log_2 \alpha_j + \log_2 X_{i,gj}$ after we ignore the small rounding error. Define $U_{i,gj} = \log_2 X_{i,gj}$ when $X_{i,gj} > 0$ and $U_{i,gj} = 0$ when $X_{i,gj} = 0$. We have $\delta_{ik,gj} \approx U_{i,gj} - U_{k,gj}$. We notice that the sample-specific noise, $\log_2 \alpha_j$, gets cancelled by the $\delta$-step. It is easy to show that $U_{i,gj}$ is a non-negative random variable with finite mean and variance. Let $M_{i,g}(\nu_{i,g}, \kappa, a) := EU_{i,gj}$ and $\sigma_{i,g}^2(\nu_{i,g}, \kappa, a) := \mathrm{var}(U_{i,gj})$, we have

$$E\delta_{ik,gj} \approx M_{i,g} - M_{k,g}, \; \mathrm{var}(\delta_{ik,gj}) \approx \sigma_{i,g}^2 + \sigma_{k,g}^2. \tag{A2.8}$$

Like the normal-based cases, when both $i, k \in \mathcal{S}^0, M_{i,g} - M_{k,g} = 0$; if $i \in \mathcal{S}^1$ and $k \in \mathcal{S}^0$, $M_{i,g} - M_{k,g} \neq$

0. Thus, $\delta_{ik,gj}$ carries the information of group mean differences encoded in $\nu_{i,g}$.

According to the Central Limit Theorem, although $\delta_{ik,gj}$ is not a normal random variable, $\bar{\delta}_{ik,g\cdot} \sim AN(M_{i,g} - M_{k,g}, \frac{\sigma_{i,g}^2+\sigma_{k,g}^2}{N_g})$ when $N_g$ is large. Similarly, we can show that $\left(\widetilde{\mathrm{Res}}_{i,gj}\right)^2 := \left(\tilde{y}_{i,gj} - \bar{\tilde{y}}_{i,g\cdot}\right)^2$, where $\tilde{y}_{i,gj} := W_{i,gj} - \bar{W}_{\cdot,gj}$ is the globally normalized log expression values, has finite mean and variance, therefore $\sum_{j=1}^{N_g}\left(\widetilde{\mathrm{Res}}_{i,gj}\right)^2$ is asymptotically normal. Note that $\widehat{\mathrm{BGRSS}}_i$ (defined by Equations (2.10) and (2.11) in the main text) is a smooth function of $\bar{\delta}_{ik,g\cdot}$, and $\widehat{\mathrm{WGRSS}}_i$ is a smooth function of $\sum_{j=1}^{N_g}\left(\widetilde{\mathrm{Res}}_{i,gj}\right)^2$, thus the `super-delta2` hypothesis test problem is asymptotically equivalent to a comparable normal-based `super-delta2` hypothesis test with suitable mean and variance.

Model (A2.6) is more complicated than Model (A2.5) because the technical noise is not modeled as a simple multiplicative constant but a latent factor. Recall that in this model, $\log_2(\nu_{i,g}) = \log_2 \alpha_j + \mu_{i,g}$, therefore $\nu_{i,g} = \alpha_j \cdot 2^{\mu_{i,g}}$. This implies that

$$Y_{i,gj}|\alpha_j \sim \mathrm{NBP}(\alpha_j \cdot 2^{\mu_{i,g}}, \kappa_i, a_i),\ E(Y_{i,gj}|\alpha_j) = \alpha_j \cdot 2^{\mu_{i,g}},\ \mathrm{var}(Y_{i,gj}|\alpha_j) = \alpha_j \cdot 2^{\mu_{i,g}} + \kappa(\alpha_j \cdot 2^{\mu_{i,g}})^a.$$

We will use the following first-order approximations for the log-counts

$$E(W_{i,gj}|\alpha_j) \approx \log_2 E(Y_{i,gj}|\alpha_j) = \log_2 \alpha_j + \mu_{i,g}.$$

$$\mathrm{var}(W_{i,gj}|\alpha_j) \approx \left(\frac{1}{\ln 2 \cdot E(Y_{i,gj}|\alpha_j)}\right)^2 \cdot \mathrm{var}(Y_{i,gj}|\alpha_j) = \frac{1}{(\ln 2)^2} \cdot \left(\frac{1}{\alpha_j \cdot 2^{\mu_{i,g}}} + \kappa(\alpha_j \cdot 2^{\mu_{i,g}})^{a-2}\right).$$

Based on the modeling assumption that $Y_{i,gj}$ and $Y_{k,gj}$ are independent conditional on $\alpha_j$, we have

$$E\delta_{ik,gj} \approx \mu_{i,g} - \mu_{k,g},\ \mathrm{var}(\delta_{ik,gj}|\alpha_j) \approx \sigma_{i,g}^2 + \sigma_{k,g}^2. \tag{A2.9}$$

Here $\sigma_{i,g}^2 := \frac{1}{(\ln 2)^2} \cdot \left(\frac{1}{\alpha_j \cdot 2^{\mu_{i,g}}} + \kappa(\alpha_j \cdot 2^{\mu_{i,g}})^{a-2}\right)$ and $\sigma_{k,g}^2 := \frac{1}{(\ln 2)^2} \cdot \left(\frac{1}{\alpha_j \cdot 2^{\mu_{k,g}}} + \kappa(\alpha_j \cdot 2^{\mu_{k,g}})^{a-2}\right)$ are the approximate conditional variances of $W_{i,gj}$ and $W_{k,gj}$, respectively. Compare Equation (A2.9) with Equation (A2.8), we find that the main technical difficulty for the latent factor model is that $\mathrm{var}(\delta_{ik,gj}|\alpha_j)$ depends on the realization of $\alpha_j$, therefore we need the following additional assumptions to ensure the marginal variance of $\delta_{ik,gj}$ is finite: (a) $E\alpha_j^{-1}$ is finite, and (b) $E\alpha_j^{a-2}$ is finite. Apparently, the uniform distribution used in Simulations 2 and 3 satisfies these two conditions.

With these two additional assumptions, we can ensure that

$$\begin{aligned}\mathrm{var}(\delta_{ik,gj}) &= E\mathrm{var}(\delta_{ik,gj}|\alpha_j) + \mathrm{var}\left(E(\delta_{ik,gj}|\alpha_j)\right) \approx E(\sigma_{i,g}^2 + \sigma_{k,g}^2) + 0 \\ &= \frac{1}{(\ln 2)^2} \cdot \left(E\alpha_j^{-1}\left(\frac{1}{2^{\mu_{i,g}}} + \frac{1}{2^{\mu_{k,g}}}\right) + \kappa E\alpha_j^{a-2}\left(2^{(a-2)\mu_{i,g}} + 2^{(a-2)\mu_{k,g}}\right)\right) < \infty.\end{aligned} \tag{A2.10}$$

With finite mean and variance, we can now use the central limit theorem to show that as $N_g \to \infty$,

$$\sqrt{N_g}\left(\bar{\delta}_{ik,g\cdot} - (\mu_{i,g} - \mu_{k,g})\right) \xrightarrow{\text{in distribution}} N\left(0, \mathrm{var}(\delta_{ik,gj})\right).$$

Therefore, the `super-delta2` F-test for this model is asymptotically equivalent to a comparable normal-based `super-delta2` hypothesis test with suitable mean and variance.

### Appendix 3: Super-delta2 Algorithm

As a summary, we have our updated `super-delta2` algorithm as follows.

1. Calculate $\widehat{\mathrm{WGRSS}}_i$ by Equations (2.5) and (2.8). Specifically:
   a. Apply global normalization to obtain $\tilde{y}_{i,gj}$.
   b. Calculate $\widetilde{\mathrm{Res}}_{i,gj}$ based on Equation (2.5),
   c. Calculate $\widehat{\mathrm{WGRSS}}_i$ and $\hat{\sigma}_i^2$ based on Equation (2.8).
2. Calculate $R_{ik,g}$ by Equations (2.9).
3. Obtain $R_{i,g}^{\mathrm{trim}}$ (Rtrim1) by applying spherical trimming to $R_{ik,g}$, and take per-group trimmed mean.
4. Apply the bias-correction described in Appendix 3 to $R_{i,g}^{\mathrm{trim}}$ to obtain trimmed and bias-corrected estimate of BGRSS, denoted as $\widehat{\mathrm{BGRSS}}_i$.
5. Calculate the F-statistics by $F_i := \frac{N-G}{G-1} \cdot \frac{\widehat{\mathrm{BGRSS}}_i}{\widehat{\mathrm{WGRSS}}_i}$ and obtain one-way ANOVA *p*-value from the F-distribution $F_{G-1,N-G}$.
6. Use *post-hoc* analysis defined by Equation (2.14) to calculate pairwise Tukey's t-statistics and their p-values for pairwise group comparisons.
7. Apply multiple testing adjustment to obtain adjusted *p*-values (for example, by Benjamini-Hochberg adjustment [2]).

### Appendix 4: The average number of differentially expressed genes selected by each method in simulation 2

| | Overall | A vs. B | A vs. C | B vs. C |
|---|---|---|---|---|
| `super-delta2` | 1023(184) | 757(211) | 598(211) | 1001(184) |
| limma+voom | 1145(400) | 860(356) | 612(255) | 1149(488) |
| edgeR | 1144(332) | 836(286) | 640(242) | 1164(420) |
| DESeq2 | 1214(392) | 874(321) | 688(282) | 1229(476) |

**Table S1**: average number of differentially expressed genes select by each method in each comparison. (·) represents the number of false positives. Due to much tighter control of type I error, `super-delta2` tended to select fewer DEGs than other methods, but it also selects significantly fewer false positives.

### Appendix 5: Boxplot of all genes in Table 6

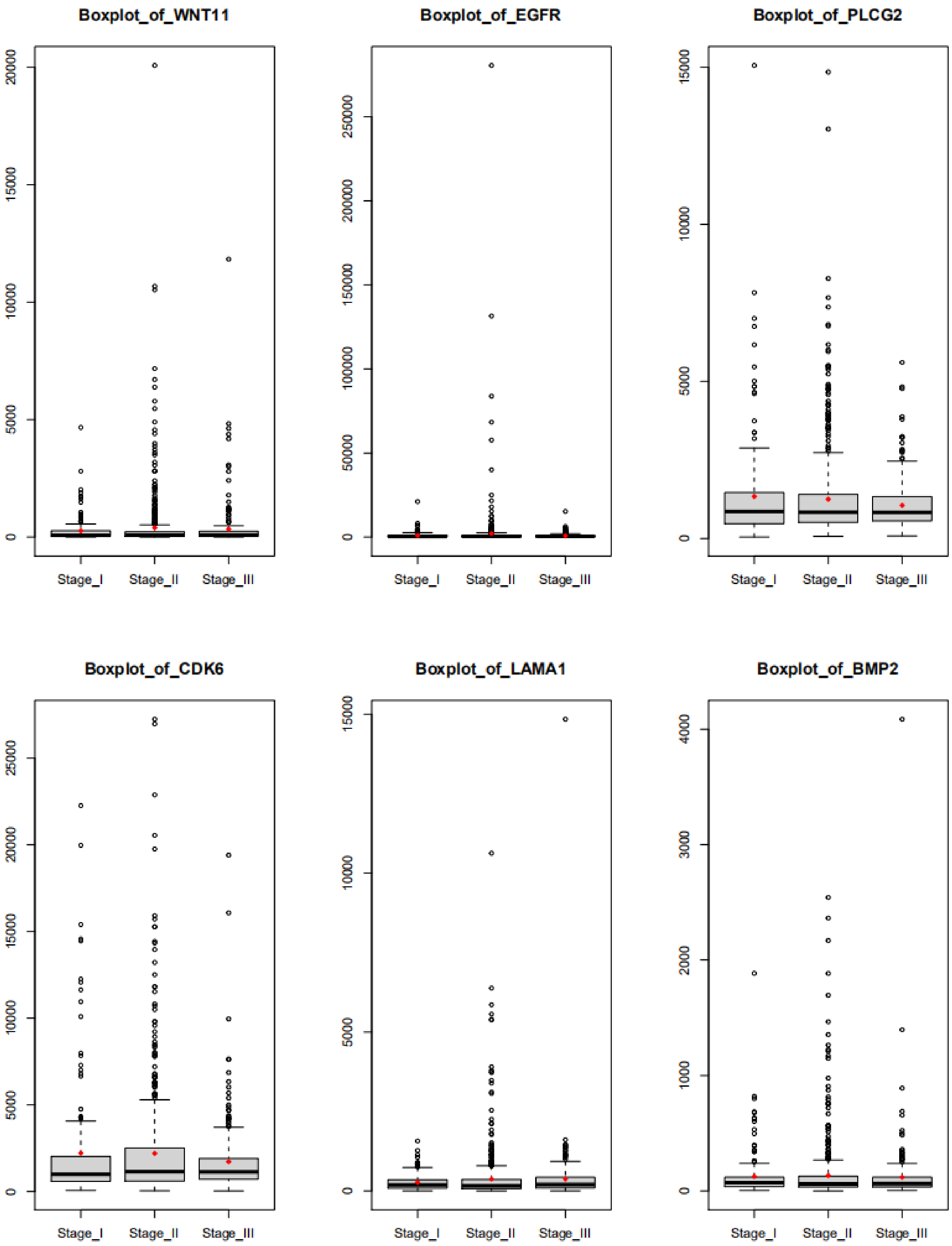


**Figure S2**: Boxplot of six genes in column 1 in Table 6.
(top six genes ranked high by DESeq2, but not `super-delta2`)

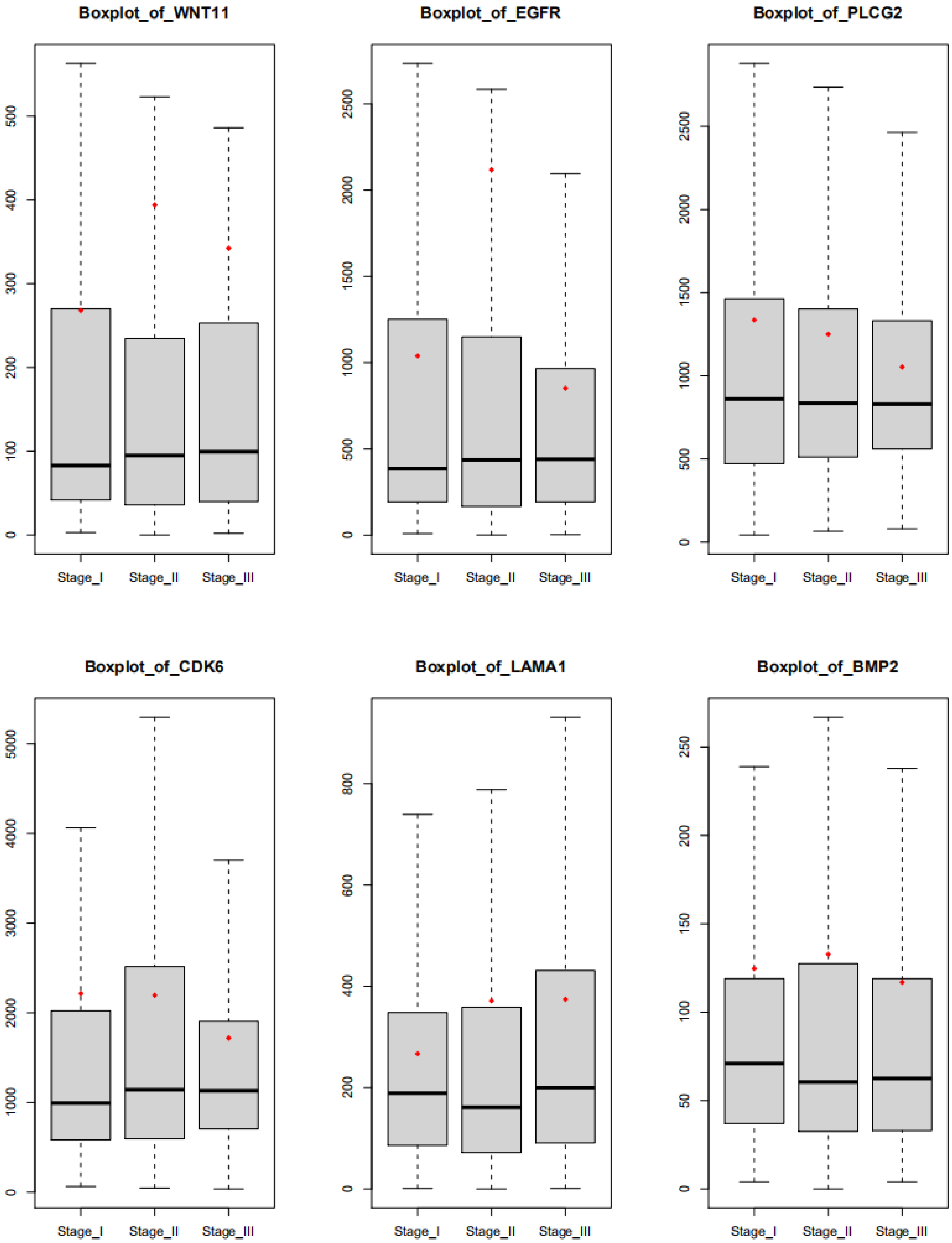


**Figure S3**: Same plots as Figure S2 without outliers.

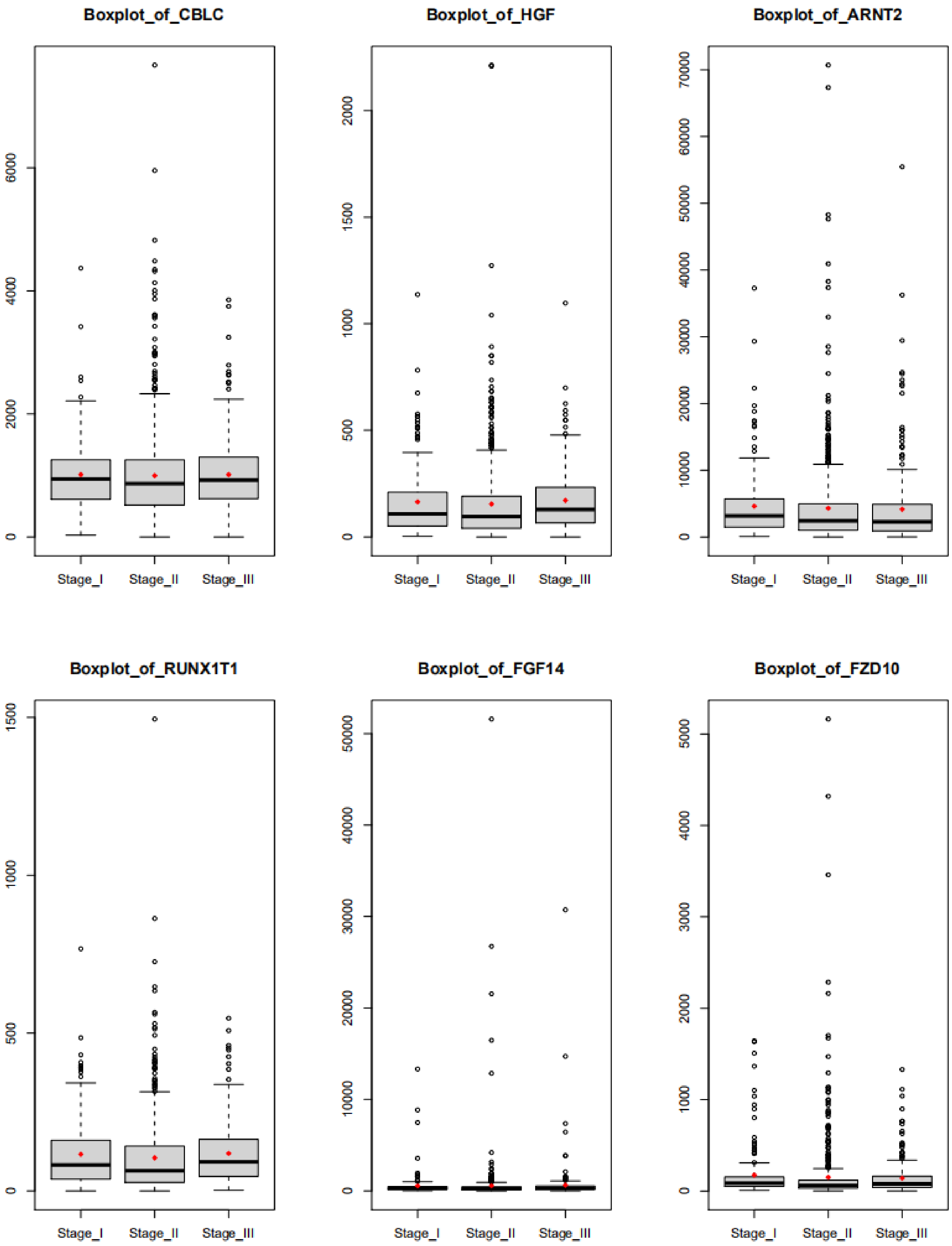


**Figure S4**: Boxplot of six genes in column 5 in Table 6.
(top 6 genes ranked high by `super-delta2`, but not DESeq2)

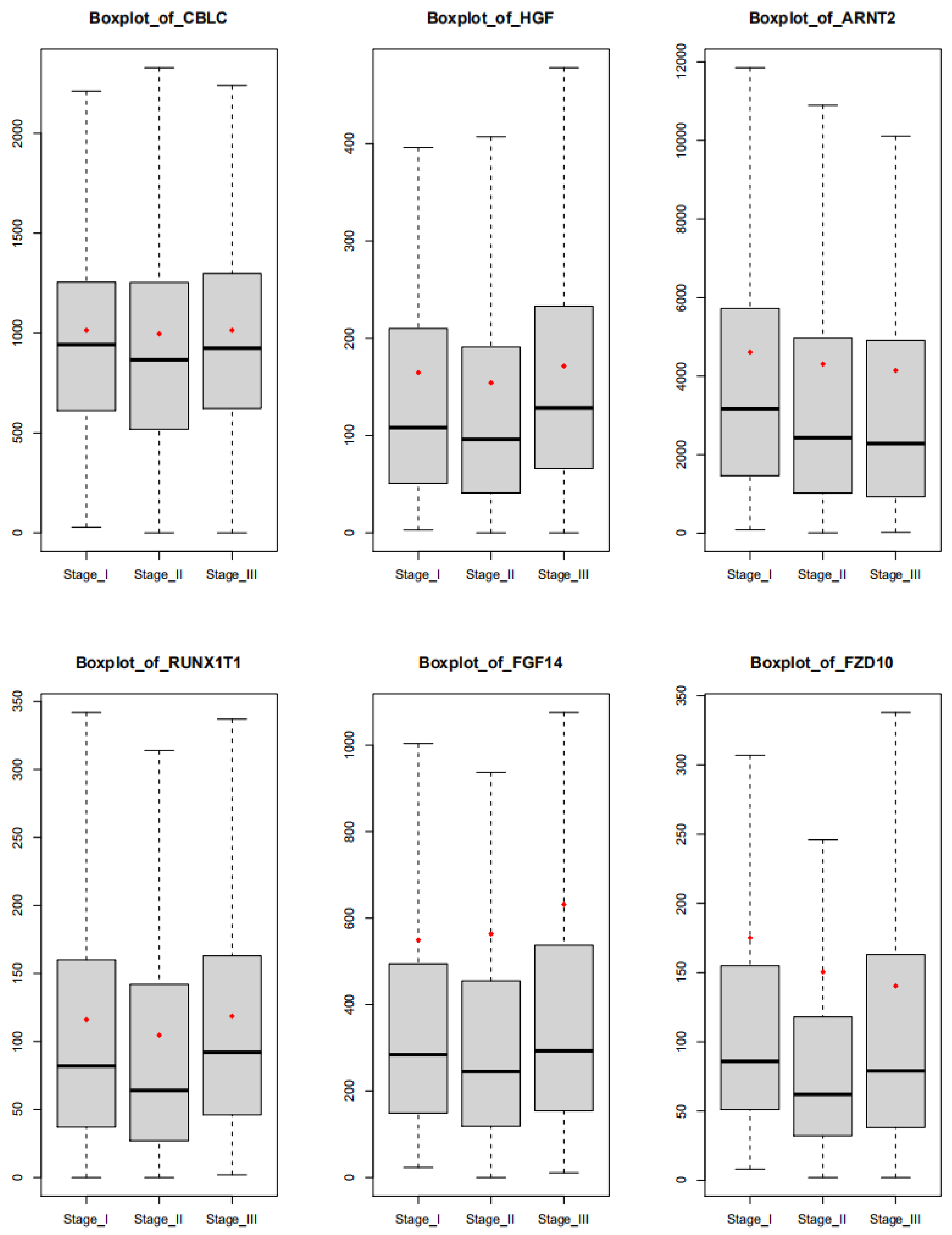


**Figure S5**: Same plots as Figure S4 without outliers.

### Appendix 6: Supplemental Simulations with Limited Sample Size

Much of the theoretical properties of `super-delta2`, when applied to RNA-seq data with Negative Binomial Poisson distribution, was developed based on normal approximation techniques in large sample theory. To understand the applicability of `super-delta2` in studies with only a few observations, we repeat the three simulations described in the main text with only $n = 10$ samples (instead of n=50 samples) in each group. To retain a reasonable level of statistical power, we compensate the greatly reduced sample size with increased effect size. Technical details of those additional simulations are listed below.

#### Supplemental Simulation 1 (SSim 1)

SSim 1 is modeled after Simulation 1 in the main text. The sample size in each group is n=10 with the following adjusted group means to increase effect size.

1. Group A was set as the baseline and $\nu_{i,g} = 100$, so that the expected mean counts of all genes are $EY_{i,gj} \approx E\alpha_{gj} \cdot \nu_{i,A} = 2100$.
2. For Group B, genes 1-600 are up-regulated with $\nu_{i,B} = 200$ so that $EY_{i,gj} \approx E\alpha_{gj} \cdot \nu_{i,B} = 4200$. Other genes have the same mean counts as group A ($\nu_{i,B} = 100,\ i = 601, \dots 5000$).
3. For Group C, genes 401-1000 are down-regulated with $\nu_{i,C} = 50$ and $EY_{i,gj} \approx 1050$. Other genes have the same mean counts as group A.

**Table S2**: Type I error rate, statistical power, and AUC value of multi-group comparisons at significance level $\alpha = 0.05$ for various methods in **SSim 1**. The 3rd column (Overall) records statistical performance of the one-way ANOVA test, the rest three columns record results from post-hoc pairwise group comparisons. All reported results are averaged over 100 repetitions. (·) represents the standard deviation of these 100 repetitions.

| | Method | Overall | A vs. B | A vs. C | B vs. C |
|---|---|---|---|---|---|
| Type I error | `super-delta2` | **0.047 (0.002)** | **0.049 (0.002)** | **0.048 (0.002)** | **0.047 (0.002)** |
| | limma+voom | 0.121 (0.007) | 0.093 (0.005) | 0.063 (0.003) | 0.153 (0.008) |
| | edgeR | 0.090 (0.004) | 0.063 (0.003) | 0.063 (0.002) | 0.114 (0.006) |
| | DESeq2 | 0.115 (0.007) | 0.076 (0.004) | 0.078 (0.004) | 0.135 (0.007) |
| Statistical power | `super-delta2` | **0.950 (0.004)** | 0.912 (0.005) | 0.930 (0.005) | **0.937 (0.005)** |
| | limma+voom | 0.900 (0.006) | 0.856 (0.007) | 0.914 (0.005) | 0.818 (0.008) |
| | edgeR | 0.940 (0.004) | 0.921 (0.005) | 0.932 (0.004) | 0.885 (0.006) |
| | DESeq2 | 0.941 (0.004) | **0.924 (0.005)** | **0.933 (0.004)** | 0.888 (0.006) |
| AUC value | `super-delta2` | **0.991 (0.001)** | **0.985 (0.002)** | **0.988 (0.001)** | **0.988 (0.002)** |
| | limma+voom | 0.961 (0.004) | 0.955 (0.004) | 0.982 (0.002) | 0.916 (0.005) |

| | | | | | |
|---|---|---|---|---|---|
| | edgeR | 0.983<br>(0.002) | 0.983<br>(0.002) | 0.985<br>(0.002) | 0.958<br>(0.004) |
| | DESeq2 | 0.977<br>(0.003) | 0.981<br>(0.002) | 0.981<br>(0.002) | 0.952<br>(0.004) |

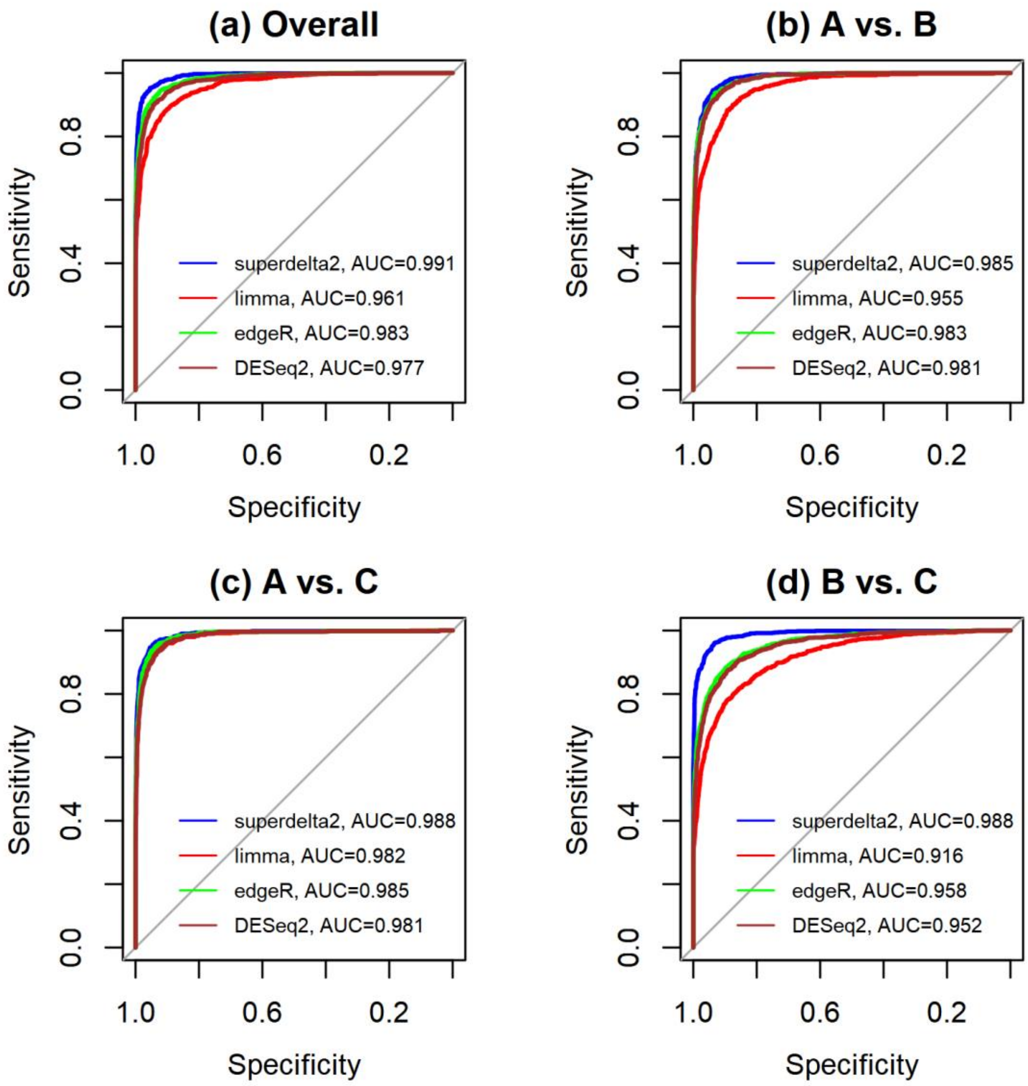


**Figure S6**: ROC curves of multi-group comparisons for SSim 1. (a) overall one-way ANOVA test; (b) Group A vs. Group B; (c) Group A vs. Group C; and (d) Group B vs. Group C.

### Supplemental Simulation 2 (SSim 2)

SSim 2 is modeled after Simulation 2 in the main text. The sample size in each group is n=10 with the following adjusted group means to increase effect size.

1. Group A was set as the baseline, where $\mu_{i,A} = \log_2 100$, so $EY_{i,gj} = 2100$.
2. For Group B, genes 1-600 are up-regulated with $\mu_{i,B} = \log_2 200$, so $EY_{i,gj} = 4200$. Other genes have the same mean counts as group A ($\nu_{i,B} = 100,\ i = 601, \dots 5000$).
3. For Group C, genes 401-1000 are down-regulated with $\mu_{i,C} = \log_2 50$, so $EY_{i,gj} = 1050$. Other genes have the same mean counts as group A.

**Table S3**: Type I error rate, statistical power, and AUC value of multi-group comparisons at significance level $\alpha = 0.05$ for various methods in **SSim 2**. The 3rd column (Overall) records statistical performance of the one-way ANOVA test, the rest three columns record results from post-hoc pairwise group comparisons. All reported results are averaged over 100 repetitions. (·) represents the standard deviation of these 100 repetitions.

| | Method | Overall | A vs. B | A vs. C | B vs. C |
|---|---|---|---|---|---|
| Type I error | `super-delta2` | **0.045 (0.002)** | **0.048 (0.002)** | **0.047 (0.002)** | **0.045 (0.003)** |
| | limma+voom | 0.090 (0.004) | 0.075 (0.004) | 0.057 (0.003) | 0.109 (0.005) |
| | edgeR | 0.071 (0.003) | 0.055 (0.003) | 0.056 (0.002) | 0.086 (0.004) |
| | DESeq2 | 0.097 (0.005) | 0.071 (0.003) | 0.074 (0.003) | 0.109 (0.005) |
| Statistical power | `super-delta2` | 0.741 (0.009) | 0.669 (0.011) | 0.704 (0.010) | **0.725 (0.010)** |
| | limma+voom | 0.678 (0.011) | 0.613 (0.012) | 0.664 (0.011) | 0.601 (0.013) |
| | edgeR | 0.767 (0.008) | 0.725 (0.009) | 0.739 (0.009) | 0.696 (0.010) |
| | DESeq2 | **0.782 (0.009)** | **0.744 (0.009)** | **0.754 (0.009)** | 0.711 (0.009) |
| AUC value | `super-delta2` | **0.946 (0.003)** | 0.914 (0.004) | **0.942 (0.003)** | **0.939 (0.003)** |
| | limma+voom | 0.891 (0.005) | 0.858 (0.007) | 0.924 (0.004) | 0.848 (0.007) |
| | edgeR | 0.935 (0.003) | **0.933 (0.003)** | 0.936 (0.003) | 0.897 (0.005) |
| | DESeq2 | 0.925 (0.003) | 0.925 (0.003) | 0.933 (0.004) | 0.891 (0.005) |

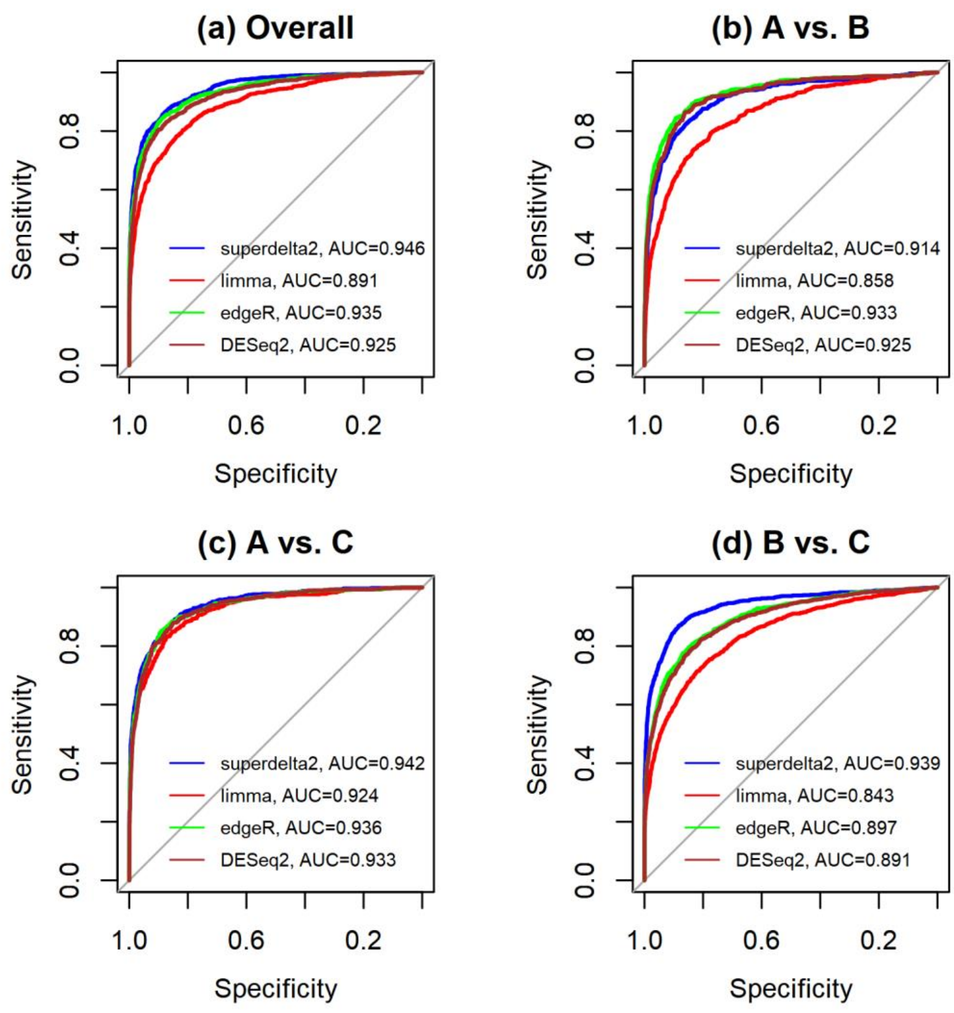


**Figure S7**: ROC curves of multi-group comparisons for SSim 2. (a) overall one-way ANOVA test; (b) Group A vs. Group B; (c) Group A vs. Group C; and (d) Group B vs. Group C.

### Supplemental Simulation 3 (SSim 3)

SSim 3 is modeled after Simulation 3 in the main text. The sample size in each group is n=10 with the following adjusted group means to increase effect size.

1. Group A was set as the baseline, where $\mu_{i,A} = \log_2 100$, so $EY_{i,gj} = 1500$.
2. For Group B, genes 1-600 are up-regulated with $\mu_{i,B} = \log_2 250$, so $EY_{i,gj} = 3750$. Other genes have the same mean counts as group A ($\nu_{i,B} = 100,\ i = 601, \dots 5000$).
3. For Group C, genes 401-1000 are down-regulated with $\mu_{i,C} = \log_2 25$, so $EY_{i,gj} = 375$. Other genes have the same mean counts as group A.

**Table S4**: Type I error rate, statistical power, and AUC value of multi-group comparisons at significance level $\alpha = 0.05$ for various methods in **SSim 3**. The 3rd column (Overall) records statistical performance of the one-way ANOVA test, the rest three columns record results from post-hoc pairwise group comparisons. All reported results are averaged over 100 repetitions. (·) represents the standard deviation of these 100 repetitions.

| | Method | Overall | A vs. B | A vs. C | B vs. C |
|---|---|---|---|---|---|
| Type I error | `super-delta2` | **0.047 (0.002)** | **0.049 (0.001)** | **0.048 (0.002)** | **0.047 (0.002)** |
| | limma+voom | 0.121 (0.006) | 0.093 (0.004) | 0.063 (0.003) | 0.153 (0.008) |
| | edgeR | 0.090 (0.004) | 0.063 (0.003) | 0.063 (0.003) | 0.114 (0.005) |
| | DESeq2 | 0.115 (0.005) | 0.076 (0.003) | 0.078 (0.003) | 0.135 (0.007) |
| Statistical power | `super-delta2` | **0.950 (0.004)** | **0.913 (0.005)** | **0.930 (0.004)** | **0.937 (0.004)** |
| | limma+voom | 0.900 (0.005) | 0.856 (0.007) | 0.914 (0.005) | 0.818 (0.009) |
| | edgeR | 0.940 (0.004) | 0.921 (0.004) | 0.932 (0.004) | 0.885 (0.005) |
| | DESeq2 | 0.941 (0.004) | 0.924 (0.004) | 0.933 (0.004) | 0.888 (0.005) |
| AUC value | `super-delta2` | **0.991 (0.001)** | **0.985 (0.001)** | **0.988 (0.001)** | **0.988 (0.001)** |
| | limma+voom | 0.961 (0.002) | 0.955 (0.003) | 0.982 (0.002) | 0.916 (0.004) |
| | edgeR | 0.983 (0.002) | 0.983 (0.002) | 0.985 (0.001) | 0.958 (0.002) |
| | DESeq2 | 0.977 (0.002) | 0.981 (0.002) | 0.981 (0.002) | 0.952 (0.003) |

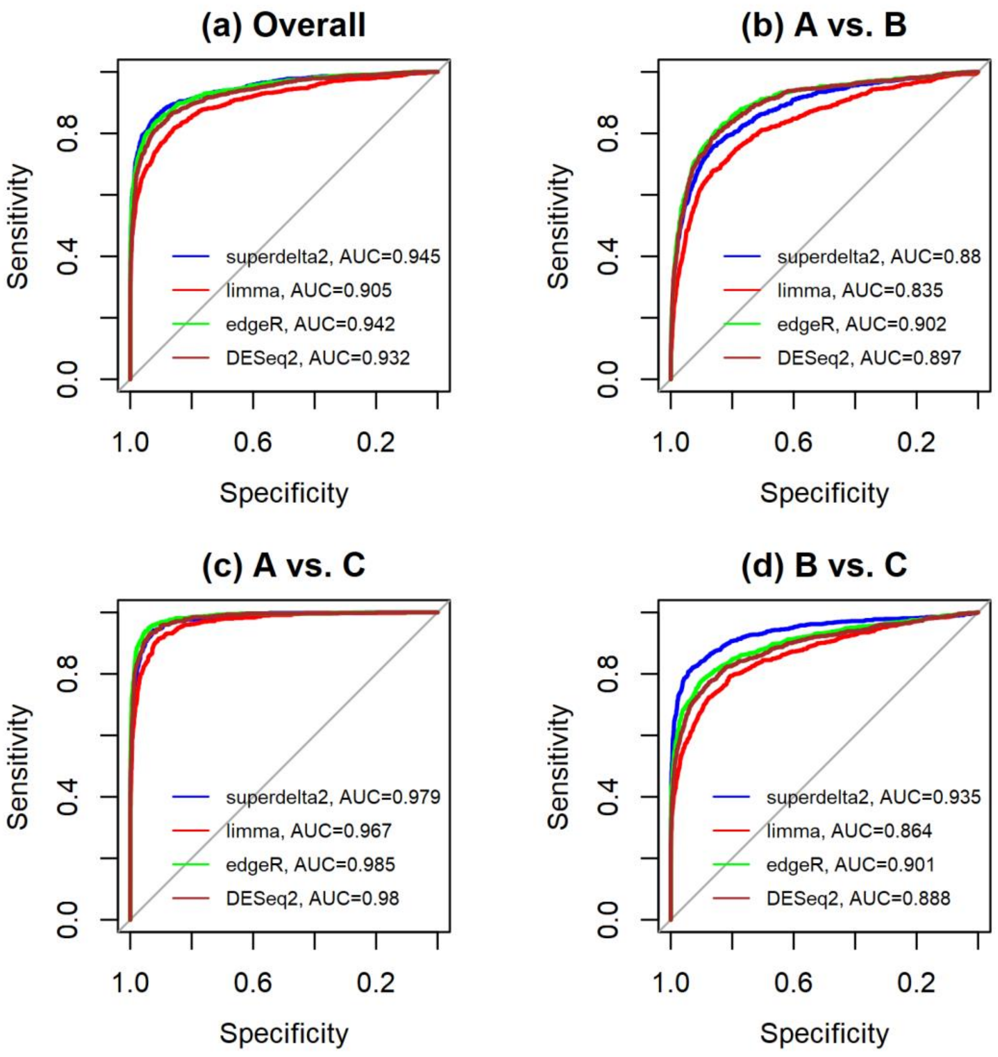


**Figure S8**: ROC curves of multi-group comparisons for SSim 3. (a) overall one-way ANOVA test; (b) Group A vs. Group B; (c) Group A vs. Group C; and (d) Group B vs. Group C.

In addition, we also tried a more extreme case with only $n = 5$ samples in each group. Unfortunately, all four methods failed to control type I error at the nominal level of $\alpha = 0.05$. For example, when we repeat SSim1 with $n = 5$, the type I errors for the overall comparison are: 0.093 (super-delta2), 0.145 (limma/voom), and 0.119 (edgeR). Of note, DESeq2 had serious convergence issues in this experimental simulation. Consequently, we were not able to obtain a reliable estimate of mean type I error for DESeq2. This poor performance is expected because: (a) the statistical models in both limma/voom and `super-delta2` rely on large sample approximations, and (b) although the statistical models used in edgeR and DESeq2 are based on GLMs therefore are exact for small samples, the hypotheses testing methods they employed (e.g., likelihood ratio test and Wald test) rely on large sample approximation, therefore are not exact for small samples either. Based on this observation, we recommend the investigators to collect at least $n = 10$ samples in each group in an RNA-seq study to avoid poor statistical performance and possible numerical issues.

### Appendix 7: Summary of the Strengths and Weaknesses of RNA-seq Differential Expression Analysis Pipelines

**Table S5**: Backgrounds, strengths, and weaknesses of the four RNA-seq differential expression analysis pipelines used in this study.

| | `super-delta2` | Limma/voom | edgeR | DESeq2 |
|---|---|---|---|---|
| Background | Based on large sample normal approximations of the NBP model. Designed for one-way multiple group comparisons | Based on large sample normal approximations of the NBP model. Designed for multiple regression problems. | Based on a generalized linear regression model with negative binomial distribution. | Based on a generalized linear regression model with negative binomial distribution, with an additional dispersion shrinkage step and a built-in outlier remover. |
| Strengths | Has a built-in trimming method to increase its robustness, controls type I error better than other methods. Computationally efficient and numerically stable. | Applicable to multivariate regression problem. Computationally efficient and numerically stable. | Applicable to multivariate regression problem. Model fitting is valid for data with small samples. Faster and numerically more robust than DESeq2. | Applicable to multivariate regression problem. Model fitting is valid for data with small samples. Its R package Provides more functionality than edgeR. |
| Weaknesses | It cannot be used for regression problems nor multi-factorial ANOVA. Its internal trimming may reduce statistical power slightly. Its inference is not exact for small sample data due to the use of large sample approximation. | It fails to control type I error at the nominal level. Its modeling fitting and statistical inference are not exact for small sample data due to the use of large sample approximation. | It requires more computational resources than `super-delta2` and limma/voom. It fails to control type I error at the nominal level. Although its model fitting is valid for small sample data, its inference is not exact due to the use of large sample approximation in p-value calculation. | It requires the most computational resources. It may fail to converge when sample size is small. It fails to control type I error at the nominal level. Although its model fitting is valid for small sample data, its inference is not exact due to the use of large sample approximation in p-value calculation. |

### Appendix 8: Comparing the Original `super-delta` method with `super-delta2`

We applied original `super-delta` method [3] on datasets we used in simulation 1 in the main text and compared the results with Table 1. Note that `super-delta` is a pairwise comparison method, so we only need to focus on the three pairwise comparisons. `Super-delta2` has a higher statistical power and both methods can control type I error well.

**Table S6**: Type I error rate, statistical power, and AUC value of multi-group comparisons at significance level $\alpha = 0.05$ for `super-delta` and `super-delta2` on simulation 1. All reported results are averaged over 100 repetitions. $(\cdot)$ represents the standard deviation of these 100 repetitions.

| | Method | A vs. B | A vs. C | B vs. C |
|---|---|---|---|---|
| Type I error | `super-delta2` | **0.050 (0.002)** | 0.048 (0.003) | **0.048 (0.002)** |
| | `super-delta` | 0.048 (0.002) | **0.049 (0.002)** | 0.048 (0.002) |
| Statistical power | `super-delta2` | **0.996 (0.001)** | **0.900 (0.006)** | **0.963 (0.004)** |
| | `super-delta` | 0.989 (0.001) | 0.882 (0.006) | 0.949 (0.005) |
| AUC value | `super-delta2` | **0.999 (0.000)** | **0.978 (0.002)** | **0.991 (0.001)** |
| | `super-delta` | 0.991 (0.001) | 0.971 (0.002) | 0.985 (0.001) |

### References


1. Di Y, Schafer DW, Cumbie JS, Chang JH: **The NBP negative binomial model for assessing differential gene expression from RNA-Seq**. *Statistical applications in genetics and molecular biology* 2011, **10**(1).
2. Benjamini Y, Hochberg Y: **Controlling the false discovery rate: a practical and powerful approach to multiple testing**. *Journal of the Royal statistical society: series B* 1995, **57**(1):289-300.
3. Liu Y, Zhang J, Qiu X: **Super-delta: a new differential gene expression analysis procedure with robust data normalization**. *BMC bioinformatics* 2017, **18**(1):582.
